# Emerin deficiency drives MCF7 cells to an invasive phenotype

**DOI:** 10.1101/2024.02.21.581379

**Authors:** Emily Hansen, Christal Rolling, Matthew Wang, James M. Holaska

## Abstract

During metastasis, cancer cells traverse the vasculature by squeezing through very small gaps in the endothelium. Thus, nuclei in metastatic cancer cells must become more malleable to move through these gaps. Our lab showed invasive breast cancer cells have 50% less emerin protein resulting in smaller, misshapen nuclei, and higher metastasis rates than non-cancerous controls. Thus, emerin deficiency was predicted to cause increased nuclear compliance, cell migration, and metastasis. We tested this hypothesis by downregulating emerin in noninvasive MCF7 cells and found emerin knockdown causes smaller, dysmorphic nuclei, resulting in increased impeded cell migration. Emerin reduction in invasive breast cancer cells showed similar results. Supporting the clinical relevance of emerin reduction in cancer progression, our analysis of 192 breast cancer patient samples showed emerin expression inversely correlates with cancer invasiveness. We conclude emerin loss is an important driver of invasive transformation and has utility as a biomarker for tumor progression.

## Introduction

Breast cancer metastasis is responsible for a majority of breast cancer-related deaths, making it a significant clinical concern.^1^ For cancer to metastasize, cancerous tumor cells must first invade the extracellular matrix and then enter the vasculature by squeezing through small gaps in the vascular endothelium. To establish a metastatic tumor, these cells travel through the body and exit the vasculature by squeezing through these gaps in the endothelium to create satellite tumors.^1^ These gaps in the endothelium are relatively small, ranging from 1.2 to 2 microns in diameter.^2^ Although the cytoplasm of cells may fit through gaps of this size, the nucleus serves as a physical barrier because it has a diameter of about 10-20 microns and a stiffness more than twice that of the cytoplasm.^3^

Nuclear morphology is well-established as an effective diagnostic tool in grading many cancers.^4^ Changes in nuclear morphology and nuclear compliance, such as nuclear softening, is associated with tumor aggressiveness and metastasis,^5–7^ and is recognized as a ‘hallmark of cancer.’^1, 8^ This nuclear softening that is associated with cancer progression allows for easier movement of cancer cells through tissues, and through the endothelial slits in the vasculature.^9^ Nuclear and cellular stiffness are also regulated by the stiffness of the tumor microenvironment (TME), caused by increased extracellular matrix (ECM) secreted by the tumor.^10^ Nuclear softness promotes invasion and metastasis, as stiffening of these invasive soft cell populations has been shown to prevent invasion in breast cancer cells.^11^

Nuclear stiffness is governed by a complex set of nucleostructural proteins that serve as signaling molecules and scaffolds. For example, emerin, an inner nuclear membrane protein that binds to lamins, is also responsible for regulating nuclear structure. ^12–14^ Interestingly, emerin is reported to be mutated in cancers, specifically in its nucleoskeletal binding domain.^13^ We previously showed that triple-negative breast cancer (TNBC) cell lines have significantly less emerin expression than their non-cancerous controls.^14^ This decreased emerin expression correlated with decreased nuclear size and increased migration and invasion.^14^ Expressing GFP-emerin rescued these deficits, while GFP-emerin mutants that failed to bind nuclear actin and lamins were unable to rescue nuclear size, migration, or invasion.^14^ In mice, we found that expressing wildtype GFP-emerin in MDA-231 cells decreased primary tumor size and lung metastasis compared to MDA-231 cells expressing GFP.^14^ GFP-emerin mutants that blocked binding to nuclear actin or lamins failed to inhibit tumor growth and metastasis in MDA-231 cells, demonstrating that emerin’s function in metastatic spread is likely dependent on its role in regulating the nucleoskeleton.^14^ On the other hand, emerin mutants that blocked emerin’s interactions with its non-structural binding partners, including barrier-to-autointegration factor (BAF) and Lim-domain only 7 (Lmo7), did rescue such phenotypes.^14^

Emerin is also implicated in prostate, hepatocellular, and lung cancers. Emerin expression inhibits metastasis in prostate cancer,^15^ supporting emerin’s involvement in metastatic disease. Conversely, without such expression of emerin, nuclei in prostate and breast cancer cell lines exhibited decreases in circularity and increases in deformity and migrasion.^15^ ^16, 17^ Hepatocellular carcinoma cells (HCC) downregulated for emerin had significantly increased cell migration and invasion compared to control HCC cells. ^16, 17^ In an additional study, decreases in emerin protein expression was also seen in 38% of ovarian cancers, and the decrease in emerin expression contributed to increased nuclear deformity marked by nuclear envelope structural defects and altered nuclear reorganization post-mitosis.^18, 19^ Interestingly, reduction of emerin protein levels was also associated with epithelial to mesenchymal transition.^20^ Thus, we predicted that emerin reduction may drive invasive phenotypes.

While these findings show that emerin has a role in metastatic disease via its nucleoskeletal interactions, they fail to ascertain whether reduced emerin protein expression is sufficient to reduce nuclear structure and increase cell migration and invasion, or if the reduced emerin expression was a result of the invasive transformation of MDA-231 cells. Thus, we tested if downregulating emerin in non-invasive cells is sufficient to convert them to a more invasive phenotype, similar to that of MDA-231 cells. Here, we show that knocking-down emerin in poorly invasive MCF7 cells does promote migration and is accompanied by smaller, more deformed nuclei, suggesting a critical link between emerin protein reduction and metastatic properties of cancer cells.

## Results

To examine the effect of emerin downregulation on MCF7, we generated MCF7 cell lines expressing either one of three different emerin shRNA sequences (A, B, C) or a scrambled shRNA sequence (Genecopoeia). We found emerin shRNA A reduced emerin protein to 39% ± 0.087 of MCF7 levels, emerin shRNA B and scrambled shRNA failed to reduce emerin expression, and emerin shRNA C reduced emerin protein expression to 42% ± 0.105 of MCF7 levels (**Figure 1**). Emerin shRNA sequence B serves as our control (now named con shRNA) because it failed to reduce emerin protein expression. Monitoring emerin mRNA by qPCR showed emerin mRNA levels are also down in emerin shRNA A, but not emerin shRNA B (Figure S1A). Thus, unless otherwise noted, the experiments using MCF7 cells were done using emerin shRNA B as the control (con shRNA), and emerin shRNA A (emerin shRNA). ^14^

**Figure 1:**
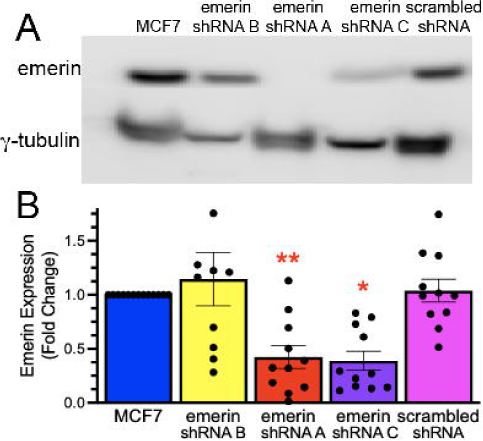
Emerin protein expression in MCF7, scrambled shRNA, and MCF7 emerin shRNA-transfected cell lines. A) Representative western blot and B) quantification of MCF7, scrambled shRNA, and three emerin shRNA cell lines normalized to γ-tubulin. *P=0.0108, **P=0.0061, N=10-13, One-way ANOVA followed by Dunnett’s multiple comparison.

To test if emerin reduction caused reduced nuclear size, we measured the nuclear area of MCF7 cells, MCF7 conshRNA cells, MCF7 scrambled shRNA cells, and MCF7 emerin shRNA cells using the ImageJ ParticleAnalyzer plug-in (see methods). Nuclear areas for MCF7 cells were 119.1 ± 2.799 µm^2^, MCF7 conshRNA was 102.6 ± 2.626 µm^2^, MCF7 scrambled shRNA was ± 106.1 ± 1.890 µm^2^, and MCF7 emerin shRNA was 78.04 ± 1.975 µm^2^. Thus, reduction of emerin protein expression caused MCF7 cells to reduce nuclear size by 35% (**Figure 2B**), while conshRNA and scrambled shRNA had minimal effects on nuclear area (14% and 11% decrease, respectively). It was possible that decreased nuclear area was caused by rounding of the nuclei, without a change in nuclear size. Thus, nuclear volume was measured in 25 or more nuclei per cell line. Nuclei from MCF7 and con shRNA cell lines were 528.9 ± 16.7 µm^3^ and 478.4 ± 18.49 µm^3^, respectively (**Figure 3A,B**). Emerin knockdown reduced nuclear volume to 369.1 ± 15.62 µm^3^ (**Figure 3A,B**,) showing emerin deficiency reduced nuclear size. To test whether nuclear structure itself was being altered, we also generated 3D renderings of nuclei from each cell line and measured nuclear curvature for easy visualization. To analyze local membrane curvature of nuclei, we used the LimeSeg plugin for Fiji.^21^

**Figure 2:**
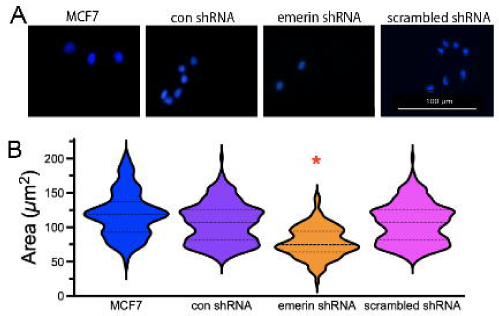
Reducing emerin in MCF7 cells decreased nuclear area. A) Representative DAPI images of MCF7, control shRNA, emerin shRNA, and scrambled shRNA cell lines, which were used to measure nuclear area. B) Violin plot of nuclear area of MCF7, control shRNA, emerin shRNA, and scrambled shRNA MCF7 cells (N>50 nuclei). The mean is depicted as the dark dashed line and the thin dashed lines represent the first and third quartiles. *P<0.0001 **P<0.0001, one-way ANOVA followed by Dunnett’s multiple comparison.

**Figure 3:**
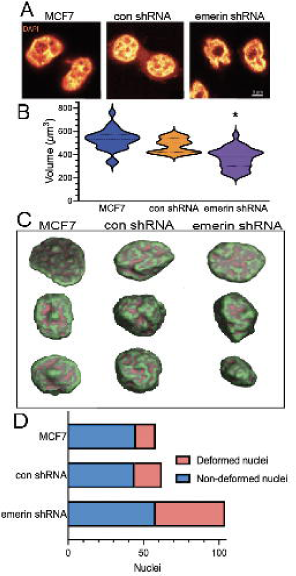
Emerin reduction decreases nuclear volume and increases nuclear bulges and indentations in MCF7 cells. A) Representative confocal images of DAPI-stained nuclei from MCF7, control shRNA, and emerin shRNA lines. B) Violin plots of nuclear volumes (N>15 nuclei for each) of MCF7, control shRNA, and emerin shRNA MCF7 cell lines. The mean is depicted as the dark dashed line and the thin dashed lines represent the first and third quartiles. *P<0.0001, one-way ANOVA followed by Dunnett’s multiple comparison test. C) 3-D rendering of z-stacks from representative nuclei in B showing the concavity and convexity of the nuclei. Red-orange indicates a concave surface and green indicates a convex surface. D) The proportion of deformed nuclei in MCF7, con shRNA, emerin shRNA, and scrambled shRNA MCF7 cells.

Using LimeSeg, we created three dimensional renders of the nuclei and then calculated the gaussian and mean curvature at 0.5 nm intervals across the surface. From this, we calculated the fraction of points that had both a positive gaussian curvature and negative mean curvature, which indicates a “concave” surface (**Figure 3C**). To measure nuclear deformity itself, nuclei were also measured, determining the proportion of nuclei with and without visible indentations and bulges. We found an increase in nuclear deformity in the emerin shRNA lines, where 44% of nuclei had visible deformity compared to the 22% deformity in MCF7 cells alone (**Figure 3D**). Thus, nuclei in the emerin-deficient MCF7 cells were smaller and more dysmorphic than controls.

Trans-well migration assays were done to test if these nuclear changes in emerin-downregulated MCF7 cells increased impeded migration. Emerin-downregulated MCF7 cells increased migration through 8 µm trans-well pores with 74.93 ± 8.131 cells/field, whereas MCF7 cells, MCF7 conshRNA cells, and MCF7 scrambled shRNA cells showed 23. ±33 ± 2.58 cells/field, 32.93 ± 2.533cells/field, and 32.93 ± 2.53 cells/field, respectively (**Figure 4A,B**). This was a 3.20-fold, 2.82-fold, and 2.28-fold increase in impeded migration when compared to MCF7, MCF7 con shRNA, and MCF7 scrambled shRNA cells, respectively (**Figure 4A,B**). To separate the nuclear structural aspects of impeded migration from the more generalized signaling and cytoskeleton reorganization associated with responding to migratory cues, we tested unimpeded migration in scratch-wound assays. However, these emerin shRNA cells did not show a difference when measuring unimpeded migration with scratch wounds (**Figure 4C,D**).

**Figure 4:**
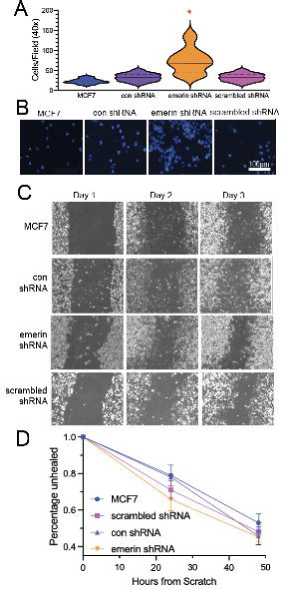
Reducing emerin in MCF7 cells increases impeded migration. A) A violin plot of the number of cells migrating through 8 µm trans-well pores is shown for MCF7, control shRNA, emerin shRNA, and scrambled shRNA cell lines (n=5 fields). The mean is depicted as the dark dashed line and the thin dashed lines represent the first and third quartiles. *P<0.0001, one-way ANOVA followed by Dunnett’s multiple comparison. C) Representative DAPI images of the cells that successfully migrated in the trans-well assays in A. C) Scratch-wound healing assay. MCF7, control shRNA, emerin shRNA, and scrambled shRNA MCF7 cell lines were plated, scratched with a pipette tip, and their migration into the wound area was monitored over three days. Representative phase images are shown. D) The rate of scratch wound healing, which refers to the ability of cells to migrate into the wound area is shown with standard error of the mean. Two-way ANOVA did not identify significance.

We then tested whether triple-negative breast cancer cells, which already express 50% less emerin compared to normal breast cells^14^ and are highly invasive, would become more invasive when emerin protein expression is reduced further. We transfected our MDA-231 cells with the same emerin shRNA plasmids as the MCF7 cells above to create stable MDA-231 cell lines. We found MDA-231 emerin shRNA lines did reduce emerin further by 50% ± 0.110 (**Figure 5B, C**) compared to MDA-231 cells. Emerin mRNA levels were also reduced compared to control shRNA and scrambled shRNA (Figure S1B). However, neither nuclear area (**Figure 5A, C-D**) nor nuclear volume (**Figure 6A-B**) decreased in emerin shRNA cell lines. As with **Figure 3**, we also tested concavity (**Figure 6C**) and deformity (**Figure 6D**) in these cells. We found an increase in nuclear deformity in the emerin shRNA lines, where 86% of nuclei had visible deformity compared to the 68% deformity in MDA-231 cells alone (**Figure 3D**). Thus, nuclei in the MDA-231 cells with further emerin-depletion were smaller and more dysmorphic than controls. This increased nuclear deformation coincided with increased migration of emerin shRNA MDA-231 cells through 8-micron trans-well pores compared to the control (**Figure 7A-B**), as emerin-downregulated MDA-231 cells increased migration through 8 µm trans-well pores with 25.67 ± 1.879 cells/field, whereas MDA-231 cells and MDA-231 conshRNA cells had 20.87 ± 1.959 cells/field and ± 25.65 ± 1.879 cells/field, respectively (**Figure 7A,B**). Downregulation of emerin had no effect on unimpeded migration (**Figure 7C-D**).

**Figure 5:**
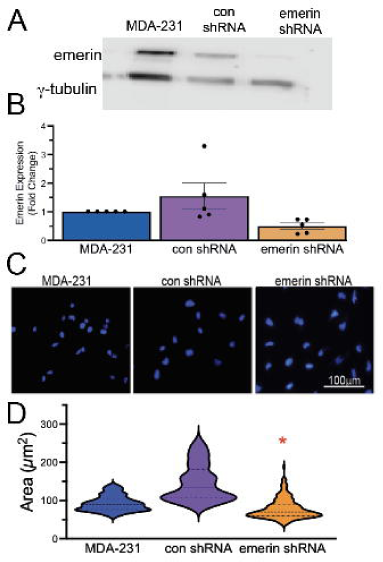
Reducing emerin in MDA-231 lines fails to affect nuclear size. A) Representative western blot of emerin and γ-tubulin (loading control) in MDA-231, control shRNA, emerin shRNA, and scrambled shRNA cell lines and B) quantitation of the western blots, N=5 (N=2 for scrambled shRNA) *P=0.0018. C) Representative images of nuclei in MDA-231, control shRNA, emerin shRNA, and scrambled shRNA stable cell lines that were used to measure nuclear area. D) A violin plot of nuclear area of MDA-231, control shRNA, emerin shRNA, and scrambled shRNA MDA-231 cell lines. The mean is depicted as the dark dashed line and the thin dashed lines represent the first and third quartiles *P<0.0006, N>50 nuclei; one-way ANOVA followed by Dunnett’s multiple comparison test.

**Figure 6:**
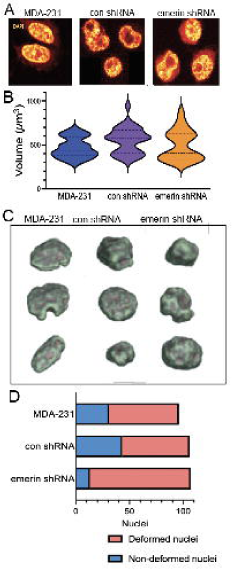
Reducing emerin in MDA-231 cells fails to decrease nuclear volume but increases nuclear deformations. A) Representative confocal images of DAPI-stained nuclei from MDA-231, control shRNA, and emerin shRNA MDA-231 cell lines. B) Violin plots of nuclear volumes (N>15 nuclei for each) of MDA-231, control shRNA, and emerin shRNA cell lines. The mean is depicted as the dark dashed line and the thin dashed lines represent the first and third quartiles. No significant differences were seen using one-way ANOVA. C) Representative images of convexity and concavity of nuclei in the respective cell lines. Red-orange indicates concave surface and green indicates a convex surface. D) Fraction of nuclei with and without deformities for MDA-231, control shRNA, and emerin shRNA cell lines.

**Figure 7:**
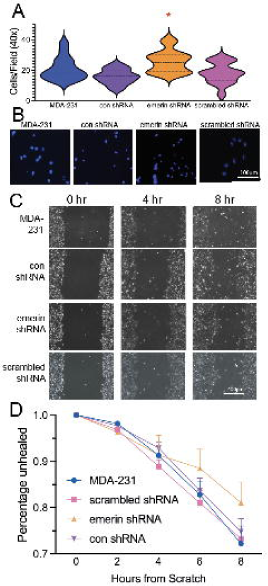
Reducing emerin in MDA-231 cells increases their impeded migration. A) A violin plot of the number of cells migrating through 8 µm trans-well pores is shown for MDA-231, control shRNA, emerin shRNA, and scrambled shRNA MDA-231 cell lines (n=5 fields). The mean is depicted as the dark dashed line and the thin dashed lines represent the first and third quartiles. *P<0.0037 compared to control shRNA, one-way ANOVA followed by Dunnett’s multiple comparison; 3 biological replicates were used for each cell line. B) Representative DAPI images of the cells that successfully migrated in the trans-well assays in A. C) Scratch-wound healing assay. MDA-231, control shRNA, emerin shRNA, and scrambled shRNA MDA-231 cell lines were plated, scratched with a pipette tip, and migration into the wound area was monitored every 2 hours for 8 hours. Representative phase images are shown. D) The rate of scratch wound healing, which refers to the ability of cells to migrate into the wound area, is shown with SEM. Two-way ANOVA was used, and no significant differences were seen.

Because emerin was implicated in regulating proliferation of MDA-231 cells,^14^ we examined cell proliferation in MCF7 cells, emerin-downregulated MCF7 cells and con shRNA MCF7 cells to test if emerin deficiency increased MCF7 proliferation. Emerin downregulation increased cell proliferation. There was a 2.5-fold increase in cell proliferation by day 6 in emerin shRNA MCF7 cells compared to the con shRNA MCF7 cells via the Presto-Blue viability assay (**Figure 8A**). Similar results were seen in MDA-231 emerin shRNA cells, in which there was a 2.5 increase in proliferation by day 6, compared to MDA-231 conshRNA cells **(Figure 8B).** It is important to note that these assays do have approximately a two-day lag time in growth before seeing such affects, which is consistent with other published cell cycle data.^22, 23^

**Figure 8:**
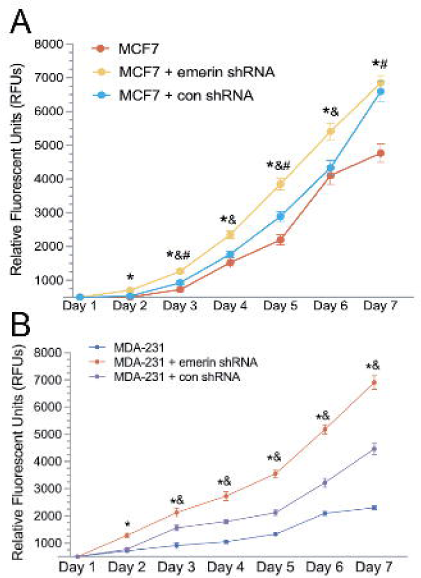
Reduction of emerin increases cell proliferation in MCF7 and MDA-231 cells. A) Growth curves of MCF7, emerin shRNA MCF7, and control shRNA MCF7 cells, as shown by measuring metabolic activity with Presto Blue Cell Viability Reagent (Life Technologies, cat#: A13261) per manufacturer’s instructions. Mean data plotted with SEM; n = 3 biological replicates. * indicates a difference between MCF7 + emerin shRNA and MCF7 cells (*p < 0.05)), as determined by two-way ANOVA and Dunnett’s test. B) Growth curves of MDA-231, emerin shRNA MDA-231, and control shRNA MDA-231 cell lines as determined using Presto Blue. Mean data plotted with SEM; n = 3 biological replicates. * indicates a significant difference between MDA-231 + emerin shRNA and MDA-231 cells (*p < 0.05), as determined by two-way ANOVA and Dunnett’s test.

These data strongly suggest that loss of emerin drives invasive cancer progression. Further, we previously found that emerin protein expression was decreased in a small sample of breast cancer patients.^14^ To rigorously determine if emerin protein expression inversely correlates with cancer invasiveness in patients, we analyzed emerin expression on 216 patient samples by immunohistochemistry with emerin antibodies (10351-1-AP, Proteintech) on a tissue microarray (TissueArray, LLC). This array contained breast cancer tumor samples from a range of types, stages, and grades (Table S1). Each tumor was assessed for emerin expression at the nuclear envelope (NE grade) based on the amount of emerin staining at the nuclear periphery; this accounts for both protein expression and normal localization.^13^ The grader was blinded to sample identifiers. After excluding tissue samples that were unable to be analyzed (i.e., damaged or folded tissues), we were able to analyze 159 samples and 12 secondary only control samples. We found emerin expression at the nuclear envelope is lower in metastatic tissue, ductal carcinoma in-situ tissue, and malignant tissue, while normal, adjacent-to-cancer normal, or benign tumor tissue had normal emerin expression at the nuclear envelope (**Figure 9A-B**). Compared to normal tissues with an average NE grade of 2.15 +/-0.1948, both invasive tissues (1.354 +/-0.0983) and metastatic tissue (0.6944 +/-0.1303) had significantly less emerin staining. To validate these results, we used a different emerin antibody from a different species (cat# 6096097, Leica) to stain an identical tissue microarray, which gave similar results in 183 samples of the same host tissue and 12 secondary control samples (**Figure 9C-D**).

**Figure 9:**
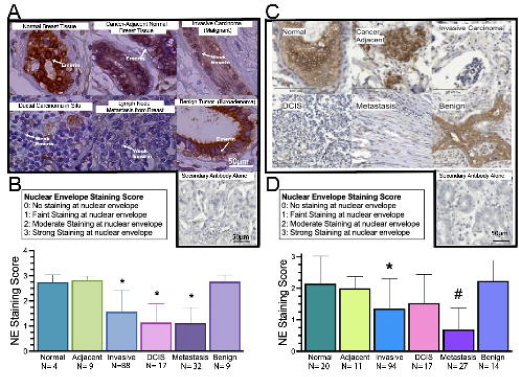
Reduced emerin expression at the nuclear periphery correlates with breast cancer invasiveness in patients. A) Representative tissue microarray staining of emerin in 159 patients using emerin polyclonal antibodies (Proteintech, cat# 10351-1-AP) or secondary alone (Vector Lab, cat#: MP-7451). Nuclei are blue, emerin is brown, and arrows denote emerin staining in certain images for reference. As severity of cases increases, there is a visible reduction in emerin expression at the nuclear envelope and more deformed nuclei are present. B) Quantification of emerin staining on IHC-stained patient samples using 0-3, with 0 having no staining at the nuclear periphery and 3 having complete, dark rim staining. N=159 total samples, *P<0.05 compared to normal tissue one-way ANOVA and Dunnett’s test. Error bars represent standard deviation. C) Representative tissue microarray staining of emerin in 183 patients using emerin monoclonal antibodies (Leica, NCL-Emerin) or secondary alone (Vector Lab, cat#: MP-7452) using the same samples in Figure 9. Nuclei are blue and emerin is brown. As aggressiveness of case increases, there is a visible reduction in emerin expression and more deformed nuclei are present. D) Quantification of emerin staining using the same grading system as in Figure 9. N=183 total samples #P<0.02 compared to all non-cancerous tissue, *P<0.0062 compared to both normal and benign tissue, one-way ANOVA and Dunnett’s test. Error bars represent standard deviation.

Normal tissues had an average NE grade of 2.82 ± 0.055 (**Figure 9A-B**). Meanwhile, metastatic tissue measured 1.128 ± 0.18 (**Figure 9A-B**) or (0.6944 +/-0.1303 (**Figure 9C-D),** as it had significantly less emerin staining. The discrepancy of tissue sample numbers is due to the quality and composition of the samples received. For example, the first trial’s samples contained a larger proportion of tissues comprised largely of connective tissue/ECM and others had folds in the tissues, leaving them unable to be included in analysis. Note that similar to the results with the emerin polyclonal antibody, emerin nuclear envelope levels were trending lower in DCIS, but more samples are needed to increase statistical power. To confirm that these results were emerin-specific, we stained this tissue microarray with just secondary antibody to account for background staining (**Figures 9A,C**).

## Discussion

It has been established for decades that the presence of abnormal nuclear structure could help distinguish tumor cells from normal cells, and to grade tumors.^24^ Yet, the contribution of these nuclear alterations to malignant transformation is unclear. Here, we show that downregulating emerin in non-invasive MCF7 cells was sufficient to decrease nuclear size, to increase nuclear deformation, and to increase impeded cell migration.

Such qualities are indicative of invasive cancers.^13, 25^ This data is consistent with previously published work, in which overexpressing emerin in TNBC lines MDA-157 and MDA-231, which have 50% less emerin than normal breast cells, increased nuclear structure and inhibited impeded migration.^13^ Importantly, the phenotypes seen by treatment with emerin shRNA were not due to the presence of the shRNA vector or cell line selection conditions because the cells containing either con shRNA or scrambled shRNA had similar phenotypes as MCF7 cells. Thus, we conclude that loss of emerin is crucial for transforming benign tumor cells to a more invasive phenotype. Consistent with our results, reduction in emerin also correlated with nuclear softening in melanoma cells,^26^ something that is well-established to correlate with rates of invasiveness and metastasis.^27, 28^

It is possible that the increased impeded migration is not due solely to changes in nuclear structure. A contribution may be caused by a disruption in mechanical signaling from the cytoplasm to the nucleus via emerin, as multiple labs have shown that disrupting the linker of nucleoskeleton and cytoskeleton (LINC) complex impairs cells’ ability to migrate.^22, 29, 30^ We predict the reduced functional interaction between emerin and the LINC complex in these cells may contribute to the migration phenotype.

Interestingly, LINC dysregulation is implicated in cancer progression and metastasis.^31–34^ In breast cancer patients, SUN1/2 and nesprins are downregulated^35^ and disruption of LINC signaling reduces nuclear F-actin,^36^ decreases nuclear size,^37^ alters MKL1 transcription,^38^ and modulates epithelial-mesenchymal transition.^39^ Since emerin binds directly to SUN1, is a major effector of LINC signaling,^40–43^ and modulates metastasis,^14^ we postulate that the loss of interaction between emerin and LINC is critical for enabling metastatic transformation Our previous work showed that the interaction of emerin with the nucleoskeleton was important for blocking metastasis.^13^ However, this study did not directly test which emerin interactions were important in the MCF7 model, so it is possible that its interactions with its other binding partners may also be playing a role. For example, emerin also binds to histone deacetylase 3 (HDAC3), Barrier-to-Autointegration Factor (BAF), and transcription factors such as Germ Cell-Less (GCL) and β-catenin, which affect the expression of their target genes.^13^ Specific to cancers, GCL has roles in regulating cell proliferation and binds the protein GAGE, which is reported to be upregulated in cancers.^44, 45^ β-catenin is also involved in proliferation, and decreasing emerin showed an increase in β-catenin and a resulting increase in proliferation,^46^ which is relevant in cancer progression. High expression of β-catenin reportedly correlates with poor patient prognosis in breast cancer.^47^ BAF is involved in DNA repair and post-translational modifications, such as recruiting chromatin regulators to the inner nuclear envelope. BAF mutations impair nuclear envelope assembly, where both emerin and lamin A are unable to assemble properly post-mitosis,^48^ which could contribute to the nuclear dysmorphism seen in cancer cells.

Knocking down emerin in MDA-231 cells did not affect nuclear area. MDA-231 cells are highly invasive TNBCs that also have 50% less emerin than normal primary breast epithelial cells, MCF10A cells,^14^ and MCF7 cells (**Figure S2**). We propose that the MDA-231 nucleus is already at is minimal size, given nucleoplasmic, chromatin, and nuclear membrane constraints, so further reduction of emerin has no effect on nuclear size. Rather emerin reduction increases the compliance of the nucleus to make it more malleable.

Consistent with these results showing emerin-deficiency drives cancer cell invasiveness, our blinded analysis of 216 breast cancer patient samples showed that lower emerin levels correlated with increased aggressiveness. These results support a model by which emerin downregulation occurs in a cell population within a growing tumor. These cells would then be selected during tumor evolution because of their increased proliferation and increased nuclear compliance, which allows them to be more invasive. This increased invasiveness enables the cells to invade the extracellular matrix and squeeze through the vascular endothelium to promote increased cancer cell survival and metastasis. Supporting our results, recent studies in prostate^15^ and ovarian cancer,^19^ found that patients have decreased emerin and that this decreased emerin contributes to higher nuclear deformity and invasion.^15^

We are aware that emerin levels do not always inversely correlate with nuclear size across all cell types.^28, 49^ However, increased nuclear deformation and increased compliance in emerin-deficient cells seem to be shared across many cell types.^49^ Whether the cell-type specificity of emerin reduction in nuclear size are caused by differences in cytoskeletal forces pushing more (to induce smaller nuclei) or less (to allow for nuclear expansion) on the nuclei, or on limiting amounts of nuclear envelope lipid components, nuclear pore complex proteins, or other nuclear envelope proteins, remains to be determined..

Collectively, these data demonstrate that emerin is critical for maintaining nuclear structure and rigidity, the loss of which makes nuclei more compliant. Based on our data, we suggest a model by which emerin expression is necessary for maintaining nuclear structure under cellular stress. Therefore, loss of emerin, such as in cancer, contributes to increased nuclear compliance, further driving tumor cell aggressiveness, invasiveness, and metastatic transformation (**Figure 10**).

**Figure 10:**
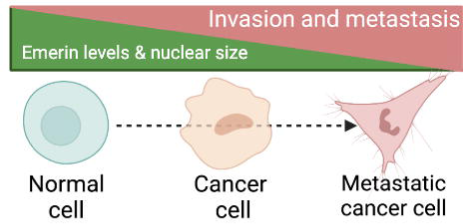
Graphical Hypothesis demonstrating the effect of emerin levels on the progression of metastatic disease.

By further investigating emerin’s role in breast cancer progression, targetable treatments for even the most invasive breast cancer types, such as triple-negative breast cancer, may be revealed. To further study this phenomenon, it will be necessary to determine how additional breast cancer subtypes respond to further emerin depletion and how depleted emerin cell lines behave in 3-D culture, which would more closely represent the 3-D tumor microenvironment. Ultimately, testing the emerin-downregulated cells used in this study in breast cancer mouse models to determine if a reduction in emerin increases tumor formation or metastasis in vivo will be needed.

However, cancer cells are not the only type of cells with low levels of emerin. For example, neutrophils also have low levels of emerin^50^ compared to other cell types, and therefore it would also be of interest to investigate this phenomenon in non-cancerous cells with reduced emerin to determine if these results are generalizable across cell types. As emerin has been shown to participate in a wide range of cellular functions,^13^ it is also possible that emerin may have different functions in different cell types, which remains to be determined.

## Materials and methods

### Cell Culture

MDA-MB-231 (ATCC cat#: HTB-26) and MCF10A (ATCC cat#: CRL-10317) were purchased from American Type Culture Collection (ATCC, Manassas, VA). MCF7 cells (ATCC cat#: HTB-22) were obtained from Mary Alpaugh’s lab (Rowan University, Camden NJ). The MDA-231 and MCF7 cells were grown in Dulbecco’s Modified Eagle Medium + GlutaMAX (Gibco cat#:10566024) with 10% Fetal Bovine Serum (Gibco, cat#: 16140089) and 1% Penicillin/Streptomycin. MCF10A cells were grown in Ham’s F-12 (Modified) + L-glutamine media (Corning, Cat#: 10-080-CV) with 5% horse serum (Gibco/Life Technologies, cat#: 16050–130), 0.5 mg/ml hydrocortisone (ThermoFisher Scientific, Waltham, MA, cat#: AC35245–0010), 100 ng/ml cholera toxin (MilliporeSigma, Burlington, MA, cat#: 227036), 10 μg/ml Insulin (Sigma-Aldrich, St. Louis, MO, cat#: 10516), and 1% Penicillin/Streptomycin. All cells were grown at 37°C and 5% CO_2_. Mycoplasma testing is done bimonthly using the MycoStrip 100 kit (InvivoGen, cat#: rep-mysnc-100).

### Creating Stable Cell Lines

All cell lines were transfected or electroporated with 2 ng/μl of each of emerin shRNA A, B, or C, (HSH095287-LVRU6MH, 3 pack, Genecopoeia) or the scrambled shRNA sequence (CSHCTR001-LVRU6MH, Genecopoeia). Specifically, scrambled shRNA and emerin shRNA C were electroporated into MCF7 cells using the Neon Electroporation System (Invitrogen). MCF7 cells were resuspended at a concentration of 2×10^5^ cells per 100 μl in buffer “R” provided with the Neon Kit, per manufacturer instructions.

Electroporation was done using 1,400 V for 10 ms for 4 pulses with the 100 μl NeonTips. Cells were plated in 6-well plates of DMEM + GlutaMAX (Gibco) + 10% FBS without penicillin/streptomycin for 24 hours before being ed with 1% penicillin/streptomycin in complete media. Lipofectamine 3000 (cat# L3000015, Invitrogen) was used to transfect emerin shRNA A and emerin shRNA B into MCF7 cells. Lipofectamine 3000 was used to transfect MDA-231 cells with scrambled shRNA, emerin shRNA A, emerin shRNA B and emerin shRNA C. For Lipofectamine 3000 transfection, cells were seeded in a 6-well plate at 70% confluency and transfected with 5 µl of Lipofectamine 3000 per reaction. All MCF7 and MDA-231 scrambled shRNA and emerin shRNA stable cell lines were selected by treating with 0.2 mg/ml hygromycin (cat#K547-20ml, VWR) after 72-hours post transfection. Cell lines were maintained at 0.8 mg/ml of hygromycin after cell line purification.

### qPCR analysis

RNA was isolated using the RNeasy^®^ Plus Mini Kit (cat # 74134, Qiagen) according to manufacturer instructions. Cells were collected from confluent 10 cm cell culture plates without selection antibiotic. qPCR was completed using Thermo Scientific Verso 1-Step qRT-PCR kit, SYBR Green, low ROX kit (cat #AB4106C, Thermo Fisher) according to manufacturer instructions, using 1 ng final concentration of the respective RNA template. qPCR primer sets for human emerin (#347828030), and human glyceraldehyde-3-phosphate dehydrogenase (GAPDH) (#345874144) were obtained from Integrated DNA Technologies (IDT) and added to the reaction at a final concentration of 1 µM. 40 qRT-PCR cycles were completed on the Quantstudio ™ 7 Pro System by Thermo Fisher Scientific.

### Immunofluorescence, volume, and nuclear area measurement

Cell lines were plated on coverslips at approximately 100,000 cells/coverslip and placed in 6-well plates. Cells were rinsed three times with 2 ml of PBS for 5 minutes and fixed with 3.7% formaldehyde in PBS for 15 minutes. Coverslips were washed again three times and permeabilized in 2 ml of 0.2% Triton X-100 in PBS for 24 minutes. Cells were blocked for 1 hour in 2 ml of 3% BSA in PBS (VWR, cat#: 97063-624). Cells were then washed three times with 2 ml of PBS for 5 minutes each and mounted with Prolong Diamond Antifade Mountant (Molecular Probes, cat#: P36971). Slides were imaged using either the Evos FL Auto Microscope using a 40x objective or Nikon confocal with 100x objective. Nuclear area was measured via Image J ParticleAnalyzer plugin. Briefly, images were transformed into a binary image where ParticleAnalyzer could then measure each nucleus. Nuclei on the edges of the image or touching other nuclei were excluded. Nuclear volume was measured via ImageJ FIJI 3D Objects Counter Plugin following the same parameters. Statistical significance was determined via one-way ANOVA followed by Dunnett’s multiple comparison test, where applicable.

### Concavity Measurements

Concavity was measured via the LimeSeg plugin for Fiji as outlined in Machado et. al.^13^ Using the plugin’s “sphere seg” command, we created three dimensional renders of the nuclei taken at 100x in a z-stack (Nikon) with 0.5-micron steps. Then, using the “ComputeCurvatures” and “DisplayCurvatures” scripts in the plugin, we calculated the mean curvature at each point along the surface at 0.5 nm intervals, and displayed regions of positive mean curvature in green and regions of negative mean curvature in red. Statistical significance was determined via one-way ANOVA followed by Dunnett’s multiple comparison test, where applicable. A minimum of 15 nuclei were rendered and measured. Deformity was determined by visually checking each nucleus at 60x magnification for regions that deviated from a spheroid or ellipsis geometry. Any nuclei found to have such deviations were counted as deformed, while nuclei that were globally convex were counted as non-deformed. The final numbers of non-deformed: deformed nuclei were determined and graphed.

### Trans-well Migration Assays

Trans-well inserts with 8-micron pores (Falcon, Cat#: 353097) were used for cell migration. Cells were plated in the chamber at a density of 1.5 ×10^5^ in serum-free media. Chambers were placed in 24-well plates containing complete growth media. After 24 hours, cells that failed to migrate through the trans-well were wiped off the top of the membrane and the cells on the bottom of the membrane were fixed for 15 minutes with 3.7% formaldehyde in PBS. The membranes were then washed 3 times in 750 µl of PBS for 5 minutes and treated with 750 µl of 0.5% Triton X-100 for 20 minutes to permeabilize membranes. The membranes were then washed a final three times with PBS and removed and mounted, cell-side down, with Prolong Diamond Antifade Mountant (Molecular Probes, cat#: P36971). Cells that migrated through were counted (five fields on the Evos FL Auto Microscope, 40x objective) after allowing the mountant to dry overnight. Statistical significance was determined via one-way ANOVA followed by Dunnett’s multiple comparison test, where applicable. At least three biological replicates were done for each cell type.

### Scratch Wound Assays

Each of the MCF7 and MDA-231 cell lines was grown to confluency in 12-well plates. A 1 mm scratch was made in each well. Plates were marked to ensure images were taken at the same location at each timepoint. Images were taken until the wound closed entirely on the Evos FL Auto Microscope using the 40x objective. Statistical significance was determined via one-way ANOVA. 3 biological replicates were done for each cell line.

### Cell Proliferation

Cell proliferation analysis was completed using the PrestoBlue Cell Viability reagent (Life Technologies, cat#: A13261) per manufacturer’s instructions. Proliferation was analyzed by plating 2 × 10^4^ cells of each cell line in a 96-well plate and cell growth was monitored every 24 hours for 7 days after plating. At least three biological replicates were done for each cell line.

### Western Blots

Whole cell lysates were suspended in NuPAGE LDS Buffer (LifeTechnologies, cat#: NP0008) with reducing agent (LifeTechnologies, cat#: NP0009). Samples were resolved by 10% SDS-PAGE gels (materials from Bio-Rad) and transferred to nitrocellulose membranes (GE Healthcare, Cat#: 10600004). Membranes were blocked in 5% nonfat dry milk (Giant brand) in PBST for two hours. Primary emerin (Protein-tech, cat# 10351-1-AP, 1:3,000 dilution) and γ-tubulin (Sigma, cat# T6557, 1:25,000 dilution) antibodies were incubated overnight with rocking at 4°C and secondary antibodies (goat anti-rabbit [cat# 31462, Invitrogen, 1:10,000] or goat anti-mouse [cat#31432, Invitrogen, 1:10,000] IgG H&L cross absorbed HRP) were incubated for 2 hours at room temperature. Emerin protein expression was normalized to γ-tubulin protein expression for quantification.

Western blots were imaged on the Li-COR Odyssey FC. Statistical significance was determined via one-way ANOVA. For **figure 10B**, protein lysates were acquired from the Debeb lab at MD Anderson Cancer Center, Houston TX. All other cell lines were grown and collected in the Holaska laboratory as described in methods.

### Immunohistochemistry and Tissue Microarray Analysis

Tissue microarrays (TissueArray, LLC, cat# BR2082c for emerin-stained samples and cat# BR087e094 for secondary-only tissue) were deparaffinized and rehydrated in coplin jars (5 minutes in xylene three times, 3 minutes in 100% EtOH three times, 3 minutes in 95% EtOH three times, 3 minutes in 80% EtOH, 3 minutes in 70% EtOH, and 5 minutes in distilled water) prior to being placed in citrate buffer (0.05% Tween 20, 10mM citric acid at pH 6.0) and steamed at 95°C for 45 minutes. Slides were cooled at RT and washed with PBS. Endogenous peroxidase was removed by washing with 0.3% hydrogen peroxide for 20 minutes at room temperature in coplin jars. After washing again with PBS, slides were blocked in 1% Bovine Serum Albumin (VWR, cat#: 97061-420) in PBS for 15 minutes. Slides were then incubated with anti-emerin antibody (Proteintech, cat#: 10351-1-AP, 1:500 dilution) for two hours at 37°C in a humidified chamber. Slides were washed again with PBS and blocked with 2.5% Normal Goat Serum for 20 minutes at room temperature (ImmPRESS Reagent, Vector Lab, cat#: MP-7451) and then incubated with the anti-rabbit ImmPRESS IgG peroxidase reagent (Vector Lab, cat#: MP-7451) or the anti-mouse ImmPRESS IgG peroxidase reagent (Vector Lab, cat# MP-7452) per manufacturer instructions. Slides were washed again with PBS and incubated with the ImmPACT DAB peroxidase substrate (Vector Lab, cat#: SK4105) for 90 seconds at room temperature while checking color development under a microscope before rinsing in tap water. Slides were counterstained with Vector Hematoxylin (Gill’s Formula, Vector Lab, cat#: H3401) for three minutes. After rinsing again in tap water, slides were incubated in 0.1% sodium bicarbonate for one minute, rinsed with distilled water, and then dehydrated and mounted with Prolong Diamond Antifade Mountant (Invitrogen, cat#: P36970). Tissue images were taken on the Evos FL Auto microscope and the Precipoint slide scanning microscope. Blinded grading of tissues was done using a grading system in which a 0 corresponded to no emerin staining at the nuclear periphery and a 3 being complete, dark staining of emerin at the nuclear periphery. All grading was done in one sitting to avoid multi-day bias. Invasive, DCIS, and metastatic tissue measurements were determined significant against noncancerous tissue *a priori* via student’s t-test.

## Acknowledgements

We thank the Department of Biomedical Sciences at Cooper Medical School of Rowan University for providing funding for this work and many fruitful discussions. We thank Dr. Isabelle Mercier (St. Joseph’s University) for many fruitful discussions regarding these studies. We thank the members of Holaska’s lab for the numerous discussions pertaining to this manuscript and the Boehning lab (Cooper Medical School at Rowan University) for technical help as needed during experiments.

## Notes

**Conflicts of Interest:** Authors have no conflicts of interest to disclose.

**Funding:** This work was supported by a grant from the National Institute of Arthritis, and Musculoskeletal and Skin Diseases (R15AR069935 to JH) and a grant from the New Jersey Commission on Cancer Research (COCR22RBG007 to JH). The content is solely the responsibility of the authors and does not necessarily represent the official views of the National Institutes of Health or the New Jersey Commission on Cancer Research. This work was also supported by Rowan University under the Camden Health Research Initiative.

### Competing Interest Statement

The authors have declared no competing interest.

### Summary of Updates

1) Changed Author order 2) Added supplemental qPCR data

## References

1. Hanahan D. Hallmarks of Cancer: New Dimensions. Cancer Discov. 2022;12(1):31–46. doi: 10.1158/2159-8290.CD-21-1059. PubMed PMID: 35022204.

2. Chaffer CL, Weinberg RA. A perspective on cancer cell metastasis. Science. 2011;331(6024):1559–64. doi: 10.1126/science.1203543. PubMed PMID: 21436443.

3. Wirtz D, Konstantopoulos K, Searson PC. The physics of cancer: the role of physical interactions and mechanical forces in metastasis. Nat Rev Cancer. 2011;11(7):512–22. Epub 20110624. doi: 10.1038/nrc3080. PubMed PMID: 21701513; PMCID: PMC3262453.

4. Bussolati G, Marchio C, Gaetano L, Lupo R, Sapino A. Pleomorphism of the nuclear envelope in breast cancer: a new approach to an old problem. J Cell Mol Med. 2008;12(1):209–18. Epub 20071205. doi: 10.1111/j.1582-4934.2007.00176.x. PubMed PMID: 18053086; PMCID: PMC3823482.

5. Acerbi I, Cassereau L, Dean I, Shi Q, Au A, Park C, Chen YY, Liphardt J, Hwang ES, Weaver VM. Human breast cancer invasion and aggression correlates with ECM stiffening and immune cell infiltration. Integr Biol (Camb). 2015;7(10):1120–34. Epub 20150511. doi: 10.1039/c5ib00040h. PubMed PMID: 25959051; PMCID: PMC4593730.

6. Kokai E, Beck H, Weissbach J, Arnold F, Sinske D, Sebert U, Gaiselmann G, Schmidt V, Walther P, Munch J, Posern G, Knoll B. Analysis of nuclear actin by overexpression of wild-type and actin mutant proteins. Histochem Cell Biol. 2014;141(2):123–35. Epub 20131004. doi: 10.1007/s00418-013-1151-4. PubMed PMID: 24091797.

7. Kai F, Laklai H, Weaver VM. Force Matters: Biomechanical Regulation of Cell Invasion and Migration in Disease. Trends Cell Biol. 2016;26(7):486–97. Epub 20160404. doi: 10.1016/j.tcb.2016.03.007. PubMed PMID: 27056543; PMCID: PMC4970516.

8. Hanahan D, Weinberg RA. Hallmarks of cancer: the next generation. Cell. 2011;144(5):646–74. doi: 10.1016/j.cell.2011.02.013. PubMed PMID: 21376230.

9. Deville SS, Cordes N. The Extracellular, Cellular, and Nuclear Stiffness, a Trinity in the Cancer Resistome-A Review. Front Oncol. 2019;9:1376. Epub 20191206. doi: 10.3389/fonc.2019.01376. PubMed PMID: 31867279; PMCID: PMC6908495.

10. Gkretsi V, Stylianopoulos T. Cell Adhesion and Matrix Stiffness: Coordinating Cancer Cell Invasion and Metastasis. Front Oncol. 2018;8:145. Epub 20180504. doi: 10.3389/fonc.2018.00145. PubMed PMID: 29780748; PMCID: PMC5945811.

11. Han YL, Pegoraro AF, Li H, Li K, Yuan Y, Xu G, Gu Z, Sun J, Hao Y, Gupta SK, Li Y, Tang W, Tang X, Teng L, Fredberg JJ, Guo M. Cell swelling, softening and invasion in a three-dimensional breast cancer model. Nat Phys. 2020;16(1):101–8. Epub 20191021. doi: 10.1038/s41567-019-0680-8. PubMed PMID: 32905405; PMCID: PMC7469976.

12. Hansen E, Holaska JM. The nuclear envelope and metastasis. Oncotarget. 2023;14:317–20. Epub 20230414. doi: 10.18632/oncotarget.28375. PubMed PMID: 37057891; PMCID: PMC10103595.

13. Liddane AG, Holaska JM. The Role of Emerin in Cancer Progression and Metastasis. Int J Mol Sci. 2021;22(20). Epub 20211019. doi: 10.3390/ijms222011289. PubMed PMID: 34681951; PMCID: PMC8537873.

14. Liddane AG, McNamara CA, Campbell MC, Mercier I, Holaska JM. Defects in Emerin-Nucleoskeleton Binding Disrupt Nuclear Structure and Promote Breast Cancer Cell Motility and Metastasis. Mol Cancer Res. 2021;19(7):1196–207. Epub 20210326. doi: 10.1158/1541-7786.MCR-20-0413. PubMed PMID: 33771882; PMCID: PMC8254762.

15. Reis-Sobreiro M, Chen JF, Novitskaya T, You S, Morley S, Steadman K, Gill NK, Eskaros A, Rotinen M, Chu CY, Chung LWK, Tanaka H, Yang W, Knudsen BS, Tseng HR, Rowat AC, Posadas EM, Zijlstra A, Di Vizio D, Freeman MR. Emerin Deregulation Links Nuclear Shape Instability to Metastatic Potential. Cancer Res. 2018;78(21):6086–97. Epub 20180828. doi: 10.1158/0008-5472.CAN-18-0608. PubMed PMID: 30154147.

16. Wu X, Wang Z, Luo L, Shu D, Wang K. Metabolomics in hepatocellular carcinoma: From biomarker discovery to precision medicine. Front Med Technol. 2022;4:1065506. Epub 20230104. doi: 10.3389/fmedt.2022.1065506. PubMed PMID: 36688143; PMCID: PMC9845953.

17. Wu KY, Xie H, Zhang ZL, Li ZX, Shi L, Zhou W, Zeng J, Tian Z, Zhang Y, Ding YB, Shen WG. Emerin knockdown induces the migration and invasion of hepatocellular carcinoma cells by up-regulating the cytoplasmic p21. Neoplasma. 2022;69(1):59–70. Epub 20211104. doi: 10.4149/neo_2021_210728N1059. PubMed PMID: 34734530.

18. Capo-chichi CD, Cai KQ, Testa JR, Godwin AK, Xu XX. Loss of GATA6 leads to nuclear deformation and aneuploidy in ovarian cancer. Mol Cell Biol. 2009;29(17):4766–77. Epub 20090706. doi: 10.1128/MCB.00087-09. PubMed PMID: 19581290; PMCID: PMC2725711.

19. Capo-chichi CD, Cai KQ, Simpkins F, Ganjei-Azar P, Godwin AK, Xu XX. Nuclear envelope structural defects cause chromosomal numerical instability and aneuploidy in ovarian cancer. BMC Med. 2011;9:28. Epub 20110326. doi: 10.1186/1741-7015-9-28. PubMed PMID: 21439080; PMCID: PMC3072346.

20. Comaills V, Kabeche L, Morris R, Buisson R, Yu M, Madden MW, LiCausi JA, Boukhali M, Tajima K, Pan S, Aceto N, Sil S, Zheng Y, Sundaresan T, Yae T, Jordan NV, Miyamoto DT, Ting DT, Ramaswamy S, Haas W, Zou L, Haber DA, Maheswaran S. Genomic Instability Is Induced by Persistent Proliferation of Cells Undergoing Epithelial-to-Mesenchymal Transition. Cell Rep. 2016;17(10):2632–47. doi: 10.1016/j.celrep.2016.11.022. PubMed PMID: 27926867; PMCID: PMC5320932.

21. Machado S, Mercier V, Chiaruttini N. LimeSeg: a coarse-grained lipid membrane simulation for 3D image segmentation. BMC Bioinformatics. 2019;20(1):2. Epub 20190103. doi: 10.1186/s12859-018-2471-0. PubMed PMID: 30606118; PMCID: PMC6318983.

22. Sutherland RL, Hall RE, Taylor IW. Cell proliferation kinetics of MCF-7 human mammary carcinoma cells in culture and effects of tamoxifen on exponentially growing and plateau-phase cells. Cancer Res. 1983;43(9):3998–4006. PubMed PMID: 6871841.

23. Cos S, Recio J, Sanchez-Barcelo EJ. Modulation of the length of the cell cycle time of MCF-7 human breast cancer cells by melatonin. Life Sci. 1996;58(9):811–6. doi: 10.1016/0024-3205(95)02359-3. PubMed PMID: 8632728.

24. Chow KH, Factor RE, Ullman KS. The nuclear envelope environment and its cancer connections. Nat Rev Cancer. 2012;12(3):196–209. Epub 20120216. doi: 10.1038/nrc3219. PubMed PMID: 22337151; PMCID: PMC4338998.

25. van Diest PJ, van der Wall E, Baak JP. Prognostic value of proliferation in invasive breast cancer: a review. J Clin Pathol. 2004;57(7):675–81. doi: 10.1136/jcp.2003.010777. PubMed PMID: 15220356; PMCID: PMC1770351.

26. Lavenus SB, Vosatka KW, Caruso AP, Ullo MF, Khan A, Logue JS. Emerin regulation of nuclear stiffness is required for fast amoeboid migration in confined environments. J Cell Sci. 2022;135(8). Epub 20220503. doi: 10.1242/jcs.259493. PubMed PMID: 35362531.

27. Guck J, Schinkinger S, Lincoln B, Wottawah F, Ebert S, Romeyke M, Lenz D, Erickson HM, Ananthakrishnan R, Mitchell D, Kas J, Ulvick S, Bilby C. Optical deformability as an inherent cell marker for testing malignant transformation and metastatic competence. Biophys J. 2005;88(5):3689–98. Epub 20050218. doi: 10.1529/biophysj.104.045476. PubMed PMID: 15722433; PMCID: PMC1305515.

28. Suresh S. Nanomedicine: elastic clues in cancer detection. Nat Nanotechnol. 2007;2(12):748–9. Epub 20071202. doi: 10.1038/nnano.2007.397. PubMed PMID: 18654425.

29. Denis KB, Cabe JI, Danielsson BE, Tieu KV, Mayer CR, Conway DE. The LINC complex is required for endothelial cell adhesion and adaptation to shear stress and cyclic stretch. Mol Biol Cell. 2021;32(18):1654–63. Epub 20210630. doi: 10.1091/mbc.E20-11-0698. PubMed PMID: 34191529; PMCID: PMC8684736.

30. Lombardi ML, Jaalouk DE, Shanahan CM, Burke B, Roux KJ, Lammerding J. The interaction between nesprins and sun proteins at the nuclear envelope is critical for force transmission between the nucleus and cytoskeleton. J Biol Chem. 2011;286(30):26743–53. Epub 20110607. doi: 10.1074/jbc.M111.233700. PubMed PMID: 21652697; PMCID: PMC3143636.

31. Sur-Erdem I, Hussain MS, Asif M, Pinarbasi N, Aksu AC, Noegel AA. Nesprin-1 impact on tumorigenic cell phenotypes. Mol Biol Rep. 2020;47(2):921–34. Epub 20191118. doi: 10.1007/s11033-019-05184-w. PubMed PMID: 31741263.

32. Lv XB, Liu L, Cheng C, Yu B, Xiong L, Hu K, Tang J, Zeng L, Sang Y. SUN2 exerts tumor suppressor functions by suppressing the Warburg effect in lung cancer. Sci Rep. 2015;5:17940. Epub 20151210. doi: 10.1038/srep17940. PubMed PMID: 26658802; PMCID: PMC4674702.

33. Chen X, Chen Y, Huang HM, Li HD, Bu FT, Pan XY, Yang Y, Li WX, Li XF, Huang C, Meng XM, Li J. SUN2: A potential therapeutic target in cancer. Oncol Lett. 2019;17(2):1401–8. Epub 20181127. doi: 10.3892/ol.2018.9764. PubMed PMID: 30675193; PMCID: PMC6341589.

34. Liu L, Li SW, Yuan W, Tang J, Sang Y. Downregulation of SUN2 promotes metastasis of colon cancer by activating BDNF/TrkB signalling by interacting with SIRT1. J Pathol. 2021;254(5):531–42. Epub 20210602. doi: 10.1002/path.5697. PubMed PMID: 33931868.

35. Matsumoto A, Hieda M, Yokoyama Y, Nishioka Y, Yoshidome K, Tsujimoto M, Matsuura N. Global loss of a nuclear lamina component, lamin A/C, and LINC complex components SUN1, SUN2, and nesprin-2 in breast cancer. Cancer Med. 2015;4(10):1547–57. Epub 20150714. doi: 10.1002/cam4.495. PubMed PMID: 26175118; PMCID: PMC4618625.

36. Plessner M, Melak M, Chinchilla P, Baarlink C, Grosse R. Nuclear F-actin formation and reorganization upon cell spreading. J Biol Chem. 2015;290(18):11209–16. Epub 20150310. doi: 10.1074/jbc.M114.627166. PubMed PMID: 25759381; PMCID: PMC4416828.

37. Porter L, Minaisah RM, Ahmed S, Ali S, Norton R, Zhang Q, Ferraro E, Molenaar C, Holt M, Cox S, Fountain S, Shanahan C, Warren D. SUN1/2 Are Essential for RhoA/ROCK-Regulated Actomyosin Activity in Isolated Vascular Smooth Muscle Cells. Cells. 2020;9(1). Epub 20200106. doi: 10.3390/cells9010132. PubMed PMID: 31935926; PMCID: PMC7017107.

38. Hu X, Liu ZZ, Chen X, Schulz VP, Kumar A, Hartman AA, Weinstein J, Johnston JF, Rodriguez EC, Eastman AE, Cheng J, Min L, Zhong M, Carroll C, Gallagher PG, Lu J, Schwartz M, King MC, Krause DS, Guo S. MKL1-actin pathway restricts chromatin accessibility and prevents mature pluripotency activation. Nat Commun. 2019;10(1):1695. Epub 20190412. doi: 10.1038/s41467-019-09636-6. PubMed PMID: 30979898; PMCID: PMC6461646.

39. Dejardin T, Carollo PS, Sipieter F, Davidson PM, Seiler C, Cuvelier D, Cadot B, Sykes C, Gomes ER, Borghi N. Nesprins are mechanotransducers that discriminate epithelial-mesenchymal transition programs. J Cell Biol. 2020;219(10). doi: 10.1083/jcb.201908036. PubMed PMID: 32790861; PMCID: PMC7659719.

40. Chang W, Folker ES, Worman HJ, Gundersen GG. Emerin organizes actin flow for nuclear movement and centrosome orientation in migrating fibroblasts. Mol Biol Cell. 2013;24(24):3869–80. Epub 20131023. doi: 10.1091/mbc.E13-06-0307. PubMed PMID: 24152738; PMCID: PMC3861083.

41. Guilluy C, Osborne LD, Van Landeghem L, Sharek L, Superfine R, Garcia-Mata R, Burridge K. Isolated nuclei adapt to force and reveal a mechanotransduction pathway in the nucleus. Nat Cell Biol. 2014;16(4):376–81. Epub 20140309. doi: 10.1038/ncb2927. PubMed PMID: 24609268; PMCID: PMC4085695.

42. Zuela N, Zwerger M, Levin T, Medalia O, Gruenbaum Y. Impaired mechanical response of an EDMD mutation leads to motility phenotypes that are repaired by loss of prenylation. J Cell Sci. 2016;129(9):1781–91. Epub 20160331. doi: 10.1242/jcs.184309. PubMed PMID: 27034135.

43. Mandigo TR, Turcich BD, Anderson AJ, Hussey MR, Folker ES. Drosophila emerins control LINC complex localization and transcription to regulate myonuclear position. J Cell Sci. 2019;132(20). Epub 20191018. doi: 10.1242/jcs.235580. PubMed PMID: 31548202.

44. Gjerstorff MF, Rosner HI, Pedersen CB, Greve KB, Schmidt S, Wilson KL, Mollenhauer J, Besir H, Poulsen FM, Mollegaard NE, Ditzel HJ. GAGE cancer-germline antigens are recruited to the nuclear envelope by germ cell-less (GCL). PLoS One. 2012;7(9):e45819. Epub 20120920. doi: 10.1371/journal.pone.0045819. PubMed PMID: 23029259; PMCID: PMC3447759.

45. Gjerstorff MF, Ditzel HJ. An overview of the GAGE cancer/testis antigen family with the inclusion of newly identified members. Tissue Antigens. 2008;71(3):187–92. Epub 20080107. doi: 10.1111/j.1399-0039.2007.00997.x. PubMed PMID: 18179644.

46. Markiewicz E, Tilgner K, Barker N, van de Wetering M, Clevers H, Dorobek M, Hausmanowa-Petrusewicz I, Ramaekers FC, Broers JL, Blankesteijn WM, Salpingidou G, Wilson RG, Ellis JA, Hutchison CJ. The inner nuclear membrane protein emerin regulates beta-catenin activity by restricting its accumulation in the nucleus. EMBO J. 2006;25(14):3275–85. Epub 20060720. doi: 10.1038/sj.emboj.7601230. PubMed PMID: 16858403; PMCID: PMC1523183.

47. Lin SY, Xia W, Wang JC, Kwong KY, Spohn B, Wen Y, Pestell RG, Hung MC. Beta-catenin, a novel prognostic marker for breast cancer: its roles in cyclin D1 expression and cancer progression. Proc Natl Acad Sci U S A. 2000;97(8):4262–6. doi: 10.1073/pnas.060025397. PubMed PMID: 10759547; PMCID: PMC18221.

48. Lee KK, Haraguchi T, Lee RS, Koujin T, Hiraoka Y, Wilson KL. Distinct functional domains in emerin bind lamin A and DNA-bridging protein BAF. J Cell Sci. 2001;114(Pt 24):4567–73. doi: 10.1242/jcs.114.24.4567. PubMed PMID: 11792821.

49. Lammerding J, Hsiao J, Schulze PC, Kozlov S, Stewart CL, Lee RT. Abnormal nuclear shape and impaired mechanotransduction in emerin-deficient cells. J Cell Biol. 2005;170(5):781–91. Epub 20050822. doi: 10.1083/jcb.200502148. PubMed PMID: 16115958; PMCID: PMC2171355.

50. Carvalho LO, Aquino EN, Neves AC, Fontes W. The Neutrophil Nucleus and Its Role in Neutrophilic Function. J Cell Biochem. 2015;116(9):1831–6. doi: 10.1002/jcb.25124. PubMed PMID: 25727365.

