## Supplemental Table 1 for "Emerin deficiency drives MCF7 cells to an invasive phenotype"

A

| TMA Grading Figure 9A-B | | | | | | | | | | | | | |
|---|---|---|---|---|---|---|---|---|---|---|---|---|---|
| Sample location | mean score of 5 | No. | Age | Organ/Anatomic | Pathology diagnosis | Grade | Stage | Type | Tissue ID. | ER | PR | HER2 | Ki67 |
| I8 | 0 | 136 | 55 | Breast | Ductal carcinoma in situ | - | 0 | Malignant | Fmg070038 | +++ | ++ | 0 | - |
| C15 | 0 | 47 | 41 | Breast | Invasive carcinoma of no special type | 2 | IIIA | Malignant | Fmg010623 | + | - | 0 | - |
| C3 | 0 | 35 | 29 | Breast | Invasive carcinoma of no special type | 1 | IIA | Malignant | Fmg010211 | +++ | + | 0 | +, 3% |
| G4 | 0 | 100 | 47 | Breast | Invasive carcinoma of no special type | 1 | IIIB | Malignant | Fmg060196 | - | - | 0 | - |
| H4 | 0 | 116 | 59 | Breast | Invasive lobular carcinoma | - | IIA | Malignant | Fmg070189 | + | + | 0 | - |
| H11 | 0 | 123 | 48 | Breast | Squamous cell carcinoma | 1 | IIIB | Malignant | Fmg021481 | - | - | 1+ | +, 15% |
| F15 | 0.05 | 95 | 55 | Breast | Invasive carcinoma of no special type | 3 | IIA | Malignant | Fmg010730 | - | - | 0 | +, 75% |
| G16 | 0.1 | 112 | 41 | Breast | Invasive lobular carcinoma | - | IIA | Malignant | Fmg020932 | ++ | ++ | 0 | +, 15% |
| I1 | 0.125 | 129 | 54 | Breast | Intraductal papillary carcinoma | - | 0 | Malignant | Fmg041049 | +++ | ++ | 0 | +, 1% |
| F5 | 0.125 | 85 | 49 | Breast | Invasive carcinoma of no special type | 3 | IIA | Malignant | Fmg140041 | - | - | 3+ | +, 10% |
| H6 | 0.125 | 118 | 50 | Breast | Invasive lobular carcinoma | - | IIB | Malignant | Fmg070246 | - | - | 0 | - |
| A11 | 0.15 | 11 | 51 | Lymph node | Metastatic carcinoma from breast | * | - | Metastasis | Fmg020326 | +++ | ++ | 0 | +, 20% |
| I13 | 0.1666666667 | 141 | 41 | Breast | Ductal carcinoma in situ | - | 0 | Malignant | Fmg041195 | - | - | 3+ | +, 3% |
| I3 | 0.1666666667 | 131 | 49 | Breast | Ductal carcinoma in situ | - | 0 | Malignant | Fmg061057 | + | + | 0 | +, 2% |
| I15 | 0.1666666667 | 143 | 47 | Breast | Ductal carcinoma in situ (sparse) | - | 0 | Malignant | Fmg130044 | - | - | 0 | - |
| G12 | 0.2 | 108 | 70 | Breast | Invasive lobular carcinoma | - | IIIB | Malignant | Fmg020931 | ++ | - | 0 | - |
| H1 | 0.2 | 113 | 19 | Breast | Invasive lobular carcinoma | * | IIB | Malignant | Fmg070179 | ++ | + | 0 | - |
| B13 | 0.2 | 29 | 45 | Lymph node | Metastatic carcinoma from breast | 2 | - | Metastasis | Fmg100246 | - | - | 1+ | +, 3% |
| A2 | 0.25 | 2 | 55 | Lymph node | Metastatic carcinoma from breast | 2 | - | Metastasis | Fmg010606 | +++ | + | 0 | +, 3% |
| C7 | 0.3 | 39 | 54 | Breast | Invasive carcinoma of no special type | 1 | IIA | Malignant | Fmg010637 | +++ | ++ | 1+ | +, 1% |
| E11 | 0.3333333333 | 75 | 45 | Breast | Invasive carcinoma of no special type | 2 | IIA | Malignant | Fmg010622 | + | ++ | 0 | +, 3% |
| A1 | 0.35 | 1 | 40 | Lymph node | Metastatic carcinoma from breast | 2 | - | Metastasis | Fmg020433 | - | - | 3+ | +, 10% |
| F1 | 0.4 | 81 | 63 | Breast | Invasive carcinoma of no special type | 1 | IIA | Malignant | Fmg020081 | +++ | +++ | 0 | +, 10% |
| B2 | 0.4 | 18 | 28 | Lymph node | Metastatic carcinoma from breast | 2 | - | Metastasis | Fmg010628 | - | - | 3+ | +, 30% |
| B1 | 0.4 | 17 | 52 | Lymph node | Metastatic carcinoma from breast of No.44 | 2 | - | Metastasis | Fmg020652 | +++ | + | 3+ | +, 15% |
| H12 | 0.4 | 124 | 37 | Breast | Squamous cell carcinoma | 2 | IIB | Malignant | Fmg030748 | - | - | 3+ | +, 25% |
| G14 | 0.45 | 110 | 42 | Breast | Invasive lobular carcinoma | - | IIB | Malignant | Fmg060798 | - | - | 0 | - |
| C4 | 0.5 | 36 | 55 | Breast | Invasive carcinoma of no special type | 2 | IIA | Malignant | Fmg010717 | +++ | + | 0 | +, 15% |
| D1 | 0.5 | 49 | 42 | Breast | Invasive carcinoma of no special type | 3 | IIA | Malignant | Fmg031247 | - | - | 0 | +, 90% |
| D6 | 0.5 | 54 | 48 | Breast | Invasive carcinoma of no special type | 2 | IIIB | Malignant | Fmg010531 | ++ | +++ | 0 | +, 3% |
| E9 | 0.5 | 73 | 50 | Breast | Invasive carcinoma of no special type | 2 | IIIA | Malignant | Fmg010629 | - | - | 3+ | - |
| H8 | 0.5 | 120 | 44 | Breast | Invasive lobular carcinoma | - | IIA | Malignant | Fmg061070 | +++ | ++ | 0 | +, 5% |
| A7 | 0.5 | 7 | 66 | Lymph node | Metastatic carcinoma from breast | * | - | Metastasis | Fmg020534 | ++ | + | 2+ | +, 15% |
| E6 | 0.55 | 70 | 35 | Breast | Invasive carcinoma of no special type | 2 | IA | Malignant | Fmg020094 | ++ | ++ | 3+ | +, 20% |
| D2 | 0.6 | 50 | 53 | Breast | Invasive carcinoma of no special type | 3 | IIA | Malignant | Fmg010479 | +++ | - | 0 | +, 50% |
| B7 | 0.6 | 23 | 54 | Lymph node | Metastatic carcinoma from breast | 2 | - | Metastasis | Fmg100105 | ++ | + | 2+ | +, 3% |
| F12 | 0.6875 | 92 | 58 | Breast | Invasive carcinoma of no special type | 3 | IIIA | Malignant | Fmg140019 | ++ | - | 0 | +, 3% |
| A16 | 0.7 | 16 | 39 | Lymph node | Metastatic carcinoma from breast | 2 | - | Metastasis | Fmg020096 | ++ | ++ | 0 | +, 5% |
| L15 | 0.75 | 191 | 41 | Breast | Adjacent normal breast tissue | * | - | NAT | Fmg031249 | ++ | ++ | 0 | +, 3% |
| I2 | 0.75 | 130 | 43 | Breast | Ductal carcinoma in situ(breast tissue) | * | 0 | Malignant | Fmg061078 | + | + | 0 | +, 1% |
| F10 | 0.75 | 90 | 37 | Breast | Invasive carcinoma of no special type | 3 | IIB | Malignant | Fmg010612 | - | - | 0 | +, 10% |
| G1 | 0.75 | 97 | 56 | Breast | Invasive carcinoma of no special type | 1 | IIB | Malignant | Fmg110095 | - | - | 0 | +, 5% |
| H5 | 0.75 | 117 | 55 | Breast | Invasive lobular carcinoma | - | IIB | Malignant | Fmg110031 | +++ | + | 2+ | +, 5% |
| A9 | 0.8 | 9 | 44 | Lymph node | Metastatic carcinoma from breast | 2 | - | Metastasis | Fmg020543 | +++ | +++ | 0 | +, 20% |

| | | | | | | | | | | | | | |
|---|---|---|---|---|---|---|---|---|---|---|---|---|---|
| B11 | 0.8 | 27 | 59 | Lymph node | Metastatic carcinoma from breast | 2 | - | Metastasis | Fmg010537 | +++ | + | 3+ | +, 20% |
| F13 | 0.85 | 93 | 62 | Breast | Invasive carcinoma of no special type | 3 | IIA | Malignant | Fmg010732 | - | - | 0 | +, 30% |
| G5 | 0.85 | 101 | 54 | Breast | Invasive carcinoma of no special type | 1 | IIB | Malignant | Fmg140303 | - | - | 3+ | +, 3% |
| B3 | 0.85 | 19 | 80 | Lymph node | Metastatic carcinoma from breast | 2 | - | Metastasis | Fmg010939 | ++ | - | 3+ | +, 3% |
| C1 | 0.9 | 33 | 46 | Breast | Invasive carcinoma of no special type | 2 | IIB | Malignant | Fmg021511 | +++ | +++ | 0 | +, 5% |
| E1 | 0.9 | 65 | 54 | Breast | Invasive carcinoma of no special type | 2 | IA | Malignant | Fmg020088 | +++ | ++ | 3+ | +, 15% |
| E2 | 0.9 | 66 | 29 | Breast | Invasive carcinoma of no special type | 2 | IIA | Malignant | Fmg140093 | - | - | 3+ | +, 15% |
| F2 | 0.9 | 82 | 65 | Breast | Invasive carcinoma of no special type | 1 | IIA | Malignant | Fmg020082 | +++ | +++ | 2+ | +, 3% |
| A5 | 0.9 | 5 | 38 | Lymph node | Metastatic carcinoma from breast | 2 | - | Metastasis | Fmg020461 | +++ | ++ | 0 | +,20% |
| F8 | 0.95 | 88 | 63 | Breast | Invasive carcinoma of no special type | 3 | IIA | Malignant | Fmg010757 | +++ | ++ | 0 | +, 30% |
| B15 | 0.95 | 31 | 60 | Lymph node | Metastatic carcinoma from breast | 2 | - | Metastasis | Fmg010607 | - | - | 3+ | +, 65% |
| I9 | 1 | 137 | 47 | Breast | Ductal carcinoma in situ （fibrous tissue） | - | 0 | Malignant | Fmg031453 | * | * | * | * |
| C2 | 1 | 34 | 59 | Breast | Invasive carcinoma of no special type | 2 | IA | Malignant | Fmg080068 | - | - | 3+ | +,5% |
| D3 | 1 | 51 | 68 | Breast | Invasive carcinoma of no special type | 2 | IIA | Malignant | Fmg010735 | +++ | +++ | 0 | +, 3% |
| B14 | 1 | 30 | 46 | Lymph node | Metastatic carcinoma from breast | 3 | - | Metastasis | Fmg060130 | - | - | 0 | +, 70% |
| I4 | 1.05 | 132 | 45 | Breast | Ductal carcinoma in situ | - | 0 | Malignant | Fmg050351 | +++ | +++ | 0 | +, 5% |
| H9 | 1.05 | 121 | 59 | Breast | Invasive lobular carcinoma | - | IIA | Malignant | Fmg140112 | + | - | 3+ | +, 10% |
| A13 | 1.05 | 13 | 53 | Lymph node | Metastatic carcinoma from breast | 2 | - | Metastasis | Fmg020153 | - | - | 3+ | +, 10% |
| B6 | 1.05 | 22 | 49 | Lymph node | Metastatic carcinoma from breast | 3 | - | Metastasis | Fmg020530 | - | - | 3+ | +, 3% |
| D5 | 1.0625 | 53 | 50 | Breast | Invasive carcinoma of no special type | 1 | IIB | Malignant | Fmg080056 | ++ | ++ | 2+ | +, 1% |
| F4 | 1.1 | 84 | 52 | Breast | Invasive carcinoma of no special type | 3 | IIB | Malignant | Fmg110052 | ++ | - | 0 | +, 5% |
| B16 | 1.1 | 32 | 52 | Lymph node | Metastatic carcinoma from breast | 3 | - | Metastasis | Fmg010536 | +++ | - | 3+ | +, 3% |
| A3 | 1.1 | 3 | 56 | Lymph node | Metastatic carcinoma from breast with necrosis | - | - | Metastasis | Fmg010544 | +++ | + | 0 | +,15% |
| I7 | 1.125 | 135 | 39 | Breast | Ductal carcinoma in situ | - | 0 | Malignant | Fmg030989 | - | - | 3+ | +, 10% |
| I11 | 1.2 | 139 | 47 | Breast | Ductal carcinoma in situ | - | 0 | Malignant | Fmg010562 | +++ | +++ | 0 | +, 5% |
| I12 | 1.2 | 140 | 48 | Breast | Ductal carcinoma in situ | - | 0 | Malignant | Fmg031185 | +++ | +++ | 0 | +, 3% |
| I6 | 1.25 | 134 | 45 | Breast | Ductal carcinoma in situ | - | 0 | Malignant | Fmg031118 | - | - | 2+ | +, 3% |
| D4 | 1.25 | 52 | 53 | Breast | Invasive carcinoma of no special type | 2 | IIA | Malignant | Fmg030554 | +++ | - | 3+ | +, 3% |
| E15 | 1.25 | 79 | 28 | Breast | Invasive carcinoma of no special type | 3 | IB | Malignant | Fmg010409 | - | - | 3+ | +, 5% |
| H7 | 1.25 | 119 | 45 | Breast | Invasive lobular carcinoma | - | IIB | Malignant | Fmg100210 | +++ | +++ | 1+ | +, 3% |
| E7 | 1.3 | 71 | 29 | Breast | Invasive carcinoma of no special type | 2 | IIA | Malignant | Fmg010729 | - | - | 3+ | +, 5% |
| A4 | 1.3 | 4 | 43 | Lymph node | Metastatic carcinoma from breast | 2 | - | Metastasis | Fmg120014 | - | - | 3+ | +,3% |
| E3 | 1.35 | 67 | 48 | Breast | Invasive carcinoma of no special type | 2 | IIB | Malignant | Fmg140095 | +++ | +++ | 0 | +, 10% |
| G6 | 1.375 | 102 | 55 | Breast | Invasive lobular carcinoma | - | IIIA | Malignant | Fmg100027 | + | - | 3+ | - |
| C10 | 1.4 | 42 | 50 | Breast | Invasive carcinoma of no special type | 2 | IIIB | Malignant | Fmg010912 | +++ | +++ | 0 | +, 5% |
| C12 | 1.4 | 44 | 52 | Breast | Invasive carcinoma of no special type | 2 | IIB | Malignant | Fmg020652 | +++ | - | 3+ | +, 20% |
| C13 | 1.4 | 45 | 47 | Breast | Invasive carcinoma of no special type | 2 | IIB | Malignant | Fmg030518 | + | + | 3+ | +, 10% |
| A10 | 1.4 | 10 | 56 | Lymph node | Metastatic carcinoma from breast | 2 | - | Metastasis | Fmg020026 | + | - | 0 | +, 15% |
| B10 | 1.4 | 26 | 49 | Lymph node | Metastatic carcinoma from breast | 2 | - | Metastasis | Fmg080061 | +++ | - | 0 | +, 65% |
| B8 | 1.4 | 24 | 62 | Lymph node | Metastatic carcinoma from breast | 3 | - | Metastasis | Fmg060484 | + | - | 0 | +, 15% |
| I14 | 1.4375 | 142 | 55 | Breast | Ductal carcinoma in situ | - | 0 | Malignant | Fmg032009 | - | - | 3+ | +, 5% |
| D8 | 1.45 | 56 | 63 | Breast | Invasive carcinoma of no special type | 1 | IIB | Malignant | Fmg140039 | +++ | - | 2+ | +, 1% |
| E8 | 1.45 | 72 | 38 | Breast | Invasive carcinoma of no special type | 2 | IIB | Malignant | Fmg010758 | - | - | 3+ | +, 5% |
| G10 | 1.45 | 106 | 54 | Breast | Invasive lobular carcinoma | - | IIB | Malignant | Fmg100044 | ++ | + | 0 | - |
| H2 | 1.45 | 114 | 40 | Breast | Invasive lobular carcinoma | - | IIB | Malignant | Fmg060878 | ++ | + | 0 | +, 3% |
| H15 | 1.5 | 127 | 36 | Breast | Ductal carcinoma in situ with invasion | * | 0 | Malignant | Fmg050788 | +++ | +++ | 0 | +, 5% |

| | | | | | | | | | | | | | |
|---|---|---|---|---|---|---|---|---|---|---|---|---|---|
| E10 | 1.5 | 74 | 55 | Breast | Invasive carcinoma of no special type | 2 | IIIB | Malignant | Fmg010908 | ++ | - | 3+ | +, 5% |
| G7 | 1.5 | 103 | 45 | Breast | Invasive lobular carcinoma | - | IIA | Malignant | Fmg120101 | +++ | +++ | 0 | +, 3% |
| A15 | 1.5 | 15 | 38 | Lymph node | Metastatic carcinoma from breast | 2 | - | Metastasis | Fmg100103 | - | - | 0 | +, 10% |
| B12 | 1.5 | 28 | 54 | Lymph node | Metastatic carcinoma from breast | 2 | - | Metastasis | Fmg100101 | ++ | - | 2+ | +, 3% |
| H16 | 1.55 | 128 | 62 | Breast | Ductal carcinoma in situ | - | 0 | Malignant | Fmg130097 | - | - | 3+ | +, 5% |
| D12 | 1.55 | 60 | 53 | Breast | Invasive carcinoma of no special type | 2 | IIA | Malignant | Fmg010551 | - | - | 3+ | +, 20% |
| D10 | 1.6 | 58 | 32 | Breast | Invasive carcinoma of no special type | 2 | IIA | Malignant | Fmg010731 | +++ | +++ | 0 | +, 5% |
| G2 | 1.6 | 98 | 48 | Breast | Invasive carcinoma of no special type | 1 | IIIA | Malignant | Fmg110129 | - | - | 3+ | - |
| A6 | 1.6 | 6 | 56 | Lymph node | Metastatic carcinoma from breast | 2 | - | Metastasis | Fmg020431 | - | - | 3+ | +, 5% |
| B5 | 1.6 | 21 | 42 | Lymph node | Metastatic carcinoma from breast | 3 | - | Metastasis | Fmg010821 | ++ | + | 0 | - |
| G9 | 1.6875 | 105 | 69 | Breast | Invasive lobular carcinoma | - | IIA | Malignant | Fmg020930 | + | - | 0 | - |
| G3 | 1.7 | 99 | 42 | Breast | Invasive carcinoma of no special type | 1 | IIB | Malignant | Fmg110138 | +++ | - | 2+ | +, 8% |
| F11 | 1.75 | 91 | 46 | Breast | Invasive carcinoma of no special type | 3 | IIIA | Malignant | Fmg140115 | - | - | 2+ | +, 3% |
| A14 | 1.75 | 14 | 42 | Lymph node | Metastatic carcinoma from breast (sparse) | - | - | Metastasis | Fmg020539 | - | - | 3+ | +, 5% |
| G15 | 1.8 | 111 | 47 | Breast | Invasive lobular carcinoma | - | IIIB | Malignant | Fmg032391 | +++ | ++ | 0 | +, 8% |
| K1 | 1.833333333 | 161 | 37 | Breast | Ductal papillomatosis with hyperplasia | - | - | Hyperplasia | Fmg010052 | ++ | + | 0 | +, 3% |
| I16 | 1.9 | 144 | 51 | Breast | Ductal carcinoma in situ | - | 0 | Malignant | Fmg010539 | +++ | +++ | 0 | - |
| C11 | 1.9 | 43 | 62 | Breast | Invasive carcinoma of no special type | 2 | IIB | Malignant | Fmg010351 | - | - | 3+ | +, 1% |
| A12 | 1.95 | 12 | 40 | Lymph node | Metastatic carcinoma from breast | 2 | - | Metastasis | Fmg120124 | +++ | ++ | 1+ | +, 10% |
| F14 | 2 | 94 | 50 | Breast | Invasive carcinoma of no special type | 3 | IIIA | Malignant | Fmg110025 | - | - | 2+ | +, 55% |
| C6 | 2 | 38 | 64 | Breast | Invasive carcinoma of no special type (sparse) | 1 | IIA | Malignant | Fmg010825 | +++ | ++ | 0 | +, 3% |
| E16 | 2.05 | 80 | 45 | Breast | Invasive carcinoma of no special type | 1 | IIA | Malignant | Fmg010760 | +++ | +++ | 0 | +, 30% |
| G13 | 2.05 | 109 | 38 | Breast | Invasive lobular carcinoma | - | IIA | Malignant | Fmg020138 | +++ | +++ | 0 | +, 10% |
| H13 | 2.05 | 125 | 44 | Breast | Squamous cell carcinoma | 2 | IB | Malignant | Fmg090046 | - | - | 0 | +, 10% |
| G11 | 2.0625 | 107 | 59 | Breast | Invasive lobular carcinoma | - | IIIB | Malignant | Fmg020022 | +++ | +++ | 0 | +, 5% |
| C16 | 2.15 | 48 | 46 | Breast | Invasive carcinoma of no special type | 2 | IIA | Malignant | Fmg010769 | + | - | 3+ | +, 15% |
| H3 | 2.15 | 115 | 52 | Breast | Invasive lobular carcinoma | - | IIB | Malignant | Fmg031144 | - | - | 0 | +, 10% |
| J3 | 2.2 | 147 | 25 | Breast | Fibroadenoma | - | - | Benign | Fmg010046 | +++ | ++ | 0 | - |
| D16 | 2.2 | 64 | 72 | Breast | Invasive carcinoma of no special type | 2 | IIB | Malignant | Fmg080070 | - | - | 3+ | +, 5% |
| A8 | 2.2 | 8 | 55 | Lymph node | Metastatic carcinoma from breast | 2 | - | Metastasis | Fmg010941 | - | - | 0 | +, 3% |
| E13 | 2.25 | 77 | 45 | Breast | Invasive carcinoma of no special type | 2 | IIA | Malignant | Fmg010811 | - | - | 0 | +, 50% |
| C5 | 2.3 | 37 | 30 | Breast | Invasive carcinoma of no special type | 2 | IIIA | Malignant | Fmg090030 | - | - | 3+ | +, 20% |
| F16 | 2.3 | 96 | 45 | Breast | Invasive carcinoma of no special type | 3 | IIB | Malignant | Fmg110098 | +++ | - | 3+ | +, 5% |
| B4 | 2.3 | 20 | 58 | Lymph node | Metastatic carcinoma from breast | 2 | - | Metastasis | Fmg010766 | +++ | +++ | 0 | +, 3% |
| I10 | 2.35 | 138 | 63 | Breast | Ductal carcinoma in situ | - | 0 | Malignant | Fmg110044 | +++ | +++ | 2+ | +, 5% |
| F6 | 2.35 | 86 | 45 | Breast | Invasive carcinoma of no special type | * | IIA | Malignant | Fmg110141 | - | - | 0 | +, 15% |
| H14 | 2.35 | 126 | 47 | Breast | Squamous cell carcinoma | 3 | IA | Malignant | Fmg010930 | - | - | 3+ | +, 5% |
| C9 | 2.4 | 41 | 67 | Breast | Invasive carcinoma of no special type | 1 | IIA | Malignant | Fmg010624 | +++ | - | 0 | +, 5% |
| D11 | 2.4 | 59 | 52 | Breast | Invasive carcinoma of no special type | 2 | IIIB | Malignant | Fmg020350 | +++ | ++ | 0 | - |
| H10 | 2.4 | 122 | 45 | Breast | Invasive lobular carcinoma | - | IIB | Malignant | Fmg100047 | +++ | - | 2+ | +, 15% |
| D14 | 2.45 | 62 | 44 | Breast | Invasive carcinoma of no special type | 2 | IA | Malignant | Fmg010733 | +++ | +++ | 0 | +, 3% |
| F3 | 2.45 | 83 | 71 | Breast | Invasive carcinoma of no special type | 3 | IIIA | Malignant | Fmg020095 | +++ | ++ | 0 | +, 5% |
| F7 | 2.45 | 87 | 38 | Breast | Invasive carcinoma of no special type | 3 | IIA | Malignant | Fmg010860 | +++ | +++ | 0 | +, 10% |
| J6 | 2.5 | 150 | 27 | Breast | Adenosis | - | - | Inflammation | Fmg010028 | ++ | ++ | 0 | +, 8% |
| L4 | 2.5 | 180 | 42 | Breast | Cancer adjacent breast tissue | * | - | AT | Fmg020150 | + | + | 0 | - |
| J1 | 2.5 | 145 | 36 | Breast | Fibroadenoma | - | - | Benign | Fmg090075 | + | ++ | 0 | - |

| | | | | | | | | | | | | | |
|---|---|---|---|---|---|---|---|---|---|---|---|---|---|
| D9 | 2.5 | 57 | 45 | Breast | Invasive carcinoma of no special type | 2 | IA | Malignant | Fmg010547 | - | - | 0 | +, 1% |
| E5 | 2.5 | 69 | 38 | Breast | Invasive carcinoma of no special type | 2 | IIA | Malignant | Fmg010910 | + | ++ | 0 | +, 10% |
| J13 | 2.5 | 157 | 48 | Breast | Mild atypical hyperplasia of duct | - | - | Hyperplasia | Fmg010508 | +++ | ++ | 0 | - |
| K7 | 2.5 | 167 | 29 | Breast | Plasma cell mastitis | - | - | Inflammation | Fmg021965 | * | * | * | * |
| I5 | 2.6 | 133 | 46 | Breast | Ductal carcinoma in situ | - | 0 | Malignant | Fmg031516 | +++ | +++ | 0 | +, 5% |
| E4 | 2.6 | 68 | 65 | Breast | Invasive carcinoma of no special type | 2 | IIIA | Malignant | Fmg010189 | - | - | 0 | +, 10% |
| B9 | 2.6 | 25 | 39 | Lymph node | Metastatic carcinoma from breast | 2 | - | Metastasis | Fmg080063 | +++ | +++ | 2+ | - |
| D7 | 2.65 | 55 | 44 | Breast | Invasive carcinoma of no special type | 1 | IIA | Malignant | Fmg010859 | +++ | +++ | 0 | +, 10% |
| F9 | 2.65 | 89 | 47 | Breast | Invasive carcinoma of no special type | 3 | IIB | Malignant | Fmg110037 | ++ | +++ | 2+ | +, 50% |
| K6 | 2.666666667 | 166 | 39 | Breast | Chronic mastitis No.135 | * | - | Inflammation | Fmg030989 | ++ | ++ | 0 | - |
| C8 | 2.7 | 40 | 30 | Breast | Invasive carcinoma of no special type | 1 | IA | Malignant | Fmg020080 | +++ | +++ | 0 | +, 3% |
| E12 | 2.7 | 76 | 43 | Breast | Invasive carcinoma of no special type | 2 | IIA | Malignant | Fmg010835 | +++ | +++ | 0 | +, 5% |
| E14 | 2.7 | 78 | 45 | Breast | Invasive carcinoma of no special type | 2 | IIIA | Malignant | Fmg130001 | +++ | + | 3+ | +, 5% |
| J10 | 2.75 | 154 | 45 | Breast | Adenosis | - | - | Inflammation | Fmg060022 | ++ | ++ | 0 | - |
| J5 | 2.75 | 149 | 44 | Breast | Adenosis | - | - | Inflammation | Fmg010032 | ++ | ++ | 0 | - |
| L8 | 2.75 | 184 | 35 | Breast | Adjacent normal breast tissue | - | - | NAT | Fmg010800 | + | + | 0 | - |
| L10 | 2.75 | 186 | 45 | Breast | Cancer adjacent breast tissue | - | - | AT | Fmg021586 | + | + | 0 | - |
| K15 | 2.75 | 175 | 49 | Breast | Cancer adjacent breast tissue (chronic mastitis) | - | - | AT | Fmg050296 | + | + | 0 | - |
| K11 | 2.75 | 171 | 44 | Breast | Chronic mastitis | - | - | Inflammation | Fmg040540 | + | + | 0 | - |
| J2 | 2.8 | 146 | 34 | Breast | Fibroadenoma | - | - | Benign | Fmg031932 | +++ | ++ | 0 | - |
| J4 | 2.8 | 148 | 23 | Breast | Fibroadenoma | - | - | Benign | Fmg010048 | +++ | ++ | 0 | - |
| J12 | 2.833333333 | 156 | 46 | Breast | Adjacent normal breast tissue (sparse) | - | - | NAT | Fmg060211 | ++ | ++ | 0 | - |
| L7 | 2.833333333 | 183 | 46 | Breast | Cancer adjacent breast tissue | - | - | AT | Fmg030942 | + | + | 0 | +, 1% |
| J9 | 2.833333333 | 153 | 19 | Breast | Fibroadenoma | - | - | benign | Fmg010034 | ++ | ++ | 0 | +, 1% |
| D13 | 2.833333333 | 61 | 42 | Breast | Invasive carcinoma of no special type | 2 | IIA | Malignant | Fmg010917 | +++ | +++ | 0 | +, 3% |
| J15 | 2.875 | 159 | 38 | Breast | Hyperplasia | - | - | Hyperplasia | Fmg060137 | ++ | ++ | 0 | - |
| J7 | 2.9 | 151 | 23 | Breast | Fibroadenoma | - | - | Benign | Fmg010045 | +++ | ++ | 0 | +, 1% |
| K8 | 3 | 168 | 42 | Breast | Acute mastitis | - | - | Inflammation | Fmg020849 | ++ | - | 0 | - |
| L11 | 3 | 187 | 55 | Breast | Adjacent normal breast tissue | - | - | NAT | Fmg130013 | + | + | 0 | - |
| K14 | 3 | 174 | 53 | Breast | Cancer adjacent breast tissue | * | - | AT | Fmg140096 | + | + | 0 | +, 3% |
| L6 | 3 | 182 | 57 | Breast | Cancer adjacent breast tissue | * | - | AT | Fmg021442 | + | + | 0 | - |
| K13 | 3 | 173 | 29 | Breast | Chronic mastitis of No.66 | - | - | Inflammation | Fmg140093 | + | + | 0 | +, 1% |
| J8 | 3 | 152 | 45 | Breast | Fibroadenoma | - | - | Benign | Fmg060126 | ++ | +++ | 0 | +, 10% |
| J14 | 3 | 158 | 82 | Breast | Hyperplasia | - | - | Hyperplasia | Fmg060093 | + | ++ | 0 | - |
| C14 | 3 | 46 | 43 | Breast | Invasive carcinoma of no special type | * | IA | Malignant | Fmg010839 | +++ | +++ | 0 | +, 3% |
| D15 | 3 | 63 | 48 | Breast | Invasive carcinoma of no special type | 2 | IIA | Malignant | Fmg110079 | +++ | - | 2+ | +, 3% |
| J16 | 3 | 160 | 40 | Breast | Mild atypical hyperplasia of duct | - | - | Hyperplasia | Fmg060063 | +++ | +++ | 0 | +, 1% |
| K5 | 3 | 165 | 39 | Breast | Plasma cell mastitis | - | - | Inflammation | Fmg010013 | ++ | ++ | 0 | - |

**B**

| TMA Grading Figure 9C-DSample location | mean score of 5 fields | No. | Age | Organ/Anatomic Site | Pathology diagnosis | Grade | Stage | Type | Tissue ID. | ER | PR | HER2 | Ki67 |
|---|---|---|---|---|---|---|---|---|---|---|---|---|---|
| I8 | 3 | 136 | 55 | Breast | Ductal carcinoma in situ | - | 0 | Malignant | Fmg070038 | +++ | ++ | 0 | - |
| C15 | 0.5 | 47 | 41 | Breast | Invasive carcinoma of no special type | 2 | IIIA | Malignant | Fmg010623 | + | - | 0 | - |
| C3 | 2.5 | 35 | 29 | Breast | Invasive carcinoma of no special type | 1 | IIA | Malignant | Fmg010211 | +++ | + | 0 | +, 3% |
| G4 | 1 | 100 | 47 | Breast | Invasive carcinoma of no special type | 1 | IIIB | Malignant | Fmg060196 | - | - | 0 | - |
| H4 | 2.5 | 116 | 59 | Breast | Invasive lobular carcinoma | - | IIA | Malignant | Fmg070189 | + | + | 0 | - |
| H11 | 2 | 123 | 48 | Breast | Squamous cell carcinoma | 1 | IIIB | Malignant | Fmg021481 | - | - | 1+ | +, 15% |
| F15 | 0 | 95 | 55 | Breast | Invasive carcinoma of no special type | 3 | IIA | Malignant | Fmg010730 | - | - | 0 | +, 75% |

| | | | | | | | | | | | | | |
|---|---|---|---|---|---|---|---|---|---|---|---|---|---|
| G16 | 0 | 112 | 41 | Breast | Invasive lobular carcinoma | - | IIA | Malignant | Fmg020932 | ++ | ++ | 0 | +, 15% |
| I1 | 1.5 | 129 | 54 | Breast | Intraductal papillary carcinoma | - | 0 | Malignant | Fmg041049 | +++ | ++ | 0 | +, 1% |
| F5 | 1.5 | 85 | 49 | Breast | Invasive carcinoma of no special type | 3 | IIA | Malignant | Fmg140041 | - | - | 3+ | +, 10% |
| H6 | 3 | 118 | 50 | Breast | Invasive lobular carcinoma | - | IIB | Malignant | Fmg070246 | - | - | 0 | - |
| A11 | 1 | 11 | 51 | Lymph node | Metastatic carcinoma from breast | * | - | Metastasis | Fmg020326 | +++ | ++ | 0 | +, 20% |
| I13 | 1 | 141 | 41 | Breast | Ductal carcinoma in situ | - | 0 | Malignant | Fmg041195 | - | - | 3+ | +, 3% |
| I3 | 1 | 131 | 49 | Breast | Ductal carcinoma in situ | - | 0 | Malignant | Fmg061057 | + | + | 0 | +, 2% |
| I15 | 2 | 143 | 47 | Breast | Ductal carcinoma in situ (sparse) | - | 0 | Malignant | Fmg130044 | - | - | 0 | - |
| G12 | 0.5 | 108 | 70 | Breast | Invasive lobular carcinoma | - | IIIB | Malignant | Fmg020931 | ++ | - | 0 | - |
| H1 | 2 | 113 | 19 | Breast | Invasive lobular carcinoma | * | IIB | Malignant | Fmg070179 | ++ | + | 0 | - |
| B13 | 0 | 29 | 45 | Lymph node | Metastatic carcinoma from breast | 2 | - | Metastasis | Fmg100246 | - | - | 1+ | +, 3% |
| A2 | 1 | 2 | 55 | Lymph node | Metastatic carcinoma from breast | 2 | - | Metastasis | Fmg010606 | +++ | + | 0 | +, 3% |
| C7 | 1 | 39 | 54 | Breast | Invasive carcinoma of no special type | 1 | IIA | Malignant | Fmg010637 | +++ | ++ | 1+ | +, 1% |
| E11 | 0.5 | 75 | 45 | Breast | Invasive carcinoma of no special type | 2 | IIA | Malignant | Fmg010622 | + | ++ | 0 | +, 3% |
| A1 | 0 | 1 | 40 | Lymph node | Metastatic carcinoma from breast | 2 | - | Metastasis | Fmg020433 | - | - | 3+ | +, 10% |
| F1 | 0 | 81 | 63 | Breast | Invasive carcinoma of no special type | 1 | IIA | Malignant | Fmg020081 | +++ | +++ | 0 | +, 10% |
| B2 | 0.25 | 18 | 28 | Lymph node | Metastatic carcinoma from breast | 2 | - | Metastasis | Fmg010628 | - | - | 3+ | +, 30% |
| B1 | 0.5 | 17 | 52 | Lymph node | Metastatic carcinoma from breast of No.44 | 2 | - | Metastasis | Fmg020652 | +++ | + | 3+ | +, 15% |
| H12 | 2.5 | 124 | 37 | Breast | Squamous cell carcinoma | 2 | IIB | Malignant | Fmg030748 | - | - | 3+ | +, 25% |
| G14 | 2 | 110 | 42 | Breast | Invasive lobular carcinoma | - | IIB | Malignant | Fmg060798 | - | - | 0 | - |
| C4 | 2 | 36 | 55 | Breast | Invasive carcinoma of no special type | 2 | IIA | Malignant | Fmg010717 | +++ | + | 0 | +, 15% |
| D1 | 1 | 49 | 42 | Breast | Invasive carcinoma of no special type | 3 | IIA | Malignant | Fmg031247 | - | - | 0 | +, 90% |
| D6 | 3 | 54 | 48 | Breast | Invasive carcinoma of no special type | 2 | IIIB | Malignant | Fmg010531 | ++ | +++ | 0 | +, 3% |
| E9 | 2 | 73 | 50 | Breast | Invasive carcinoma of no special type | 2 | IIIA | Malignant | Fmg010629 | - | - | 3+ | - |
| H8 | 2 | 120 | 44 | Breast | Invasive lobular carcinoma | - | IIA | Malignant | Fmg061070 | +++ | ++ | 0 | +, 5% |
| A7 | 1 | 7 | 66 | Lymph node | Metastatic carcinoma from breast | * | - | Metastasis | Fmg020534 | ++ | + | 2+ | +, 15% |
| E6 | 0.5 | 70 | 35 | Breast | Invasive carcinoma of no special type | 2 | IA | Malignant | Fmg020094 | ++ | ++ | 3+ | +, 20% |
| D2 | 1.75 | 50 | 53 | Breast | Invasive carcinoma of no special type | 3 | IIA | Malignant | Fmg010479 | +++ | - | 0 | +, 50% |
| B7 | 0.5 | 23 | 54 | Lymph node | Metastatic carcinoma from breast | 2 | - | Metastasis | Fmg100105 | ++ | + | 2+ | +, 3% |
| F12 | 2 | 92 | 58 | Breast | Invasive carcinoma of no special type | 3 | IIIA | Malignant | Fmg140019 | ++ | - | 0 | +, 3% |
| A16 | | 16 | 39 | Lymph node | Metastatic carcinoma from breast | 2 | - | Metastasis | Fmg020096 | ++ | ++ | 0 | +, 5% |
| L15 | 2.5 | 191 | 41 | Breast | Adjacent normal breast tissue | * | - | NAT | Fmg031249 | ++ | ++ | 0 | +, 3% |
| I2 | 1 | 130 | 43 | Breast | Ductal carcinoma in situ(breast tissue) | * | 0 | Malignant | Fmg061078 | + | + | 0 | +, 1% |
| F10 | 0.25 | 90 | 37 | Breast | Invasive carcinoma of no special type | 3 | IIB | Malignant | Fmg010612 | - | - | 0 | +, 10% |
| G1 | 2.75 | 97 | 56 | Breast | Invasive carcinoma of no special type | 1 | IIB | Malignant | Fmg110095 | - | - | 0 | +, 5% |
| H5 | 1.5 | 117 | 55 | Breast | Invasive lobular carcinoma | - | IIB | Malignant | Fmg110031 | +++ | + | 2+ | +, 5% |
| A9 | 2 | 9 | 44 | Lymph node | Metastatic carcinoma from breast | 2 | - | Metastasis | Fmg020543 | +++ | +++ | 0 | +, 20% |
| B11 | 2 | 27 | 59 | Lymph node | Metastatic carcinoma from breast | 2 | - | Metastasis | Fmg010537 | +++ | + | 3+ | +, 20% |
| F13 | 0 | 93 | 62 | Breast | Invasive carcinoma of no special type | 3 | IIA | Malignant | Fmg010732 | - | - | 0 | +, 30% |
| G5 | 1 | 101 | 54 | Breast | Invasive carcinoma of no special type | 1 | IIB | Malignant | Fmg140303 | - | - | 3+ | +, 3% |
| B3 | 1.75 | 19 | 80 | Lymph node | Metastatic carcinoma from breast | 2 | - | Metastasis | Fmg010939 | ++ | - | 3+ | +, 3% |
| C1 | 3 | 33 | 46 | Breast | Invasive carcinoma of no special type | 2 | IIB | Malignant | Fmg021511 | +++ | +++ | 0 | +, 5% |
| E1 | 1 | 65 | 54 | Breast | Invasive carcinoma of no special type | 2 | IA | Malignant | Fmg020088 | +++ | ++ | 3+ | +, 15% |
| E2 | 1.5 | 66 | 29 | Breast | Invasive carcinoma of no special type | 2 | IIA | Malignant | Fmg140093 | - | - | 3+ | +, 15% |
| F2 | 2 | 82 | 65 | Breast | Invasive carcinoma of no special type | 1 | IIA | Malignant | Fmg020082 | +++ | +++ | 2+ | +, 3% |
| A5 | | 5 | 38 | Lymph node | Metastatic carcinoma from breast | 2 | - | Metastasis | Fmg020461 | +++ | ++ | 0 | +,20% |

| | | | | | | | | | | | | | |
|---|---|---|---|---|---|---|---|---|---|---|---|---|---|
| F8 | 1 | 88 | 63 | Breast | Invasive carcinoma of no special type | 3 | IIA | Malignant | Fmg010757 | +++ | ++ | 0 | +, 30% |
| B15 | 0 | 31 | 60 | Lymph node | Metastatic carcinoma from breast | 2 | - | Metastasis | Fmg010607 | - | - | 3+ | +, 65% |
| I9 | 2 | 137 | 47 | Breast | Ductal carcinoma in situ（fibrous tissue） | - | 0 | Malignant | Fmg031453 | * | * | * | * |
| C2 | 3 | 34 | 59 | Breast | Invasive carcinoma of no special type | 2 | IA | Malignant | Fmg080068 | - | - | 3+ | +,5% |
| D3 | 1.75 | 51 | 68 | Breast | Invasive carcinoma of no special type | 2 | IIA | Malignant | Fmg010735 | +++ | +++ | 0 | +, 3% |
| B14 | 0.5 | 30 | 46 | Lymph node | Metastatic carcinoma from breast | 3 | - | Metastasis | Fmg060130 | - | - | 0 | +, 70% |
| I4 | 2 | 132 | 45 | Breast | Ductal carcinoma in situ | - | 0 | Malignant | Fmg050351 | +++ | +++ | 0 | +, 5% |
| H9 | 0 | 121 | 59 | Breast | Invasive lobular carcinoma | - | IIA | Malignant | Fmg140112 | + | - | 3+ | +, 10% |
| A13 | 0.5 | 13 | 53 | Lymph node | Metastatic carcinoma from breast | 2 | - | Metastasis | Fmg020153 | - | - | 3+ | +, 10% |
| B6 | 1 | 22 | 49 | Lymph node | Metastatic carcinoma from breast | 3 | - | Metastasis | Fmg020530 | - | - | 3+ | +, 3% |
| D5 | 3 | 53 | 50 | Breast | Invasive carcinoma of no special type | 1 | IIB | Malignant | Fmg080056 | ++ | ++ | 2+ | +, 1% |
| F4 | 0 | 84 | 52 | Breast | Invasive carcinoma of no special type | 3 | IIB | Malignant | Fmg110052 | ++ | - | 0 | +, 5% |
| B16 | 0 | 32 | 52 | Lymph node | Metastatic carcinoma from breast | 3 | - | Metastasis | Fmg010536 | +++ | - | 3+ | +, 3% |
| A3 | 1 | 3 | 56 | Lymph node | Metastatic carcinoma from breast with necrosis | - | - | Metastasis | Fmg010544 | +++ | + | 0 | +,15% |
| I7 | 2 | 135 | 39 | Breast | Ductal carcinoma in situ | - | 0 | Malignant | Fmg030989 | - | - | 3+ | +, 10% |
| I11 | 3 | 139 | 47 | Breast | Ductal carcinoma in situ | - | 0 | Malignant | Fmg010562 | +++ | +++ | 0 | +, 5% |
| I12 | 1.5 | 140 | 48 | Breast | Ductal carcinoma in situ | - | 0 | Malignant | Fmg031185 | +++ | +++ | 0 | +, 3% |
| I6 | 2.5 | 134 | 45 | Breast | Ductal carcinoma in situ | - | 0 | Malignant | Fmg031118 | - | - | 2+ | +, 3% |
| D4 | 0 | 52 | 53 | Breast | Invasive carcinoma of no special type | 2 | IIA | Malignant | Fmg030554 | +++ | - | 3+ | +, 3% |
| E15 | 1 | 79 | 28 | Breast | Invasive carcinoma of no special type | 3 | IB | Malignant | Fmg010409 | - | - | 3+ | +, 5% |
| H7 | 0.5 | 119 | 45 | Breast | Invasive lobular carcinoma | - | IIB | Malignant | Fmg100210 | +++ | +++ | 1+ | +, 3% |
| E7 | 3 | 71 | 29 | Breast | Invasive carcinoma of no special type | 2 | IIA | Malignant | Fmg010729 | - | - | 3+ | +, 5% |
| A4 | 0 | 4 | 43 | Lymph node | Metastatic carcinoma from breast | 2 | - | Metastasis | Fmg120014 | - | - | 3+ | +,3% |
| E3 | 2.5 | 67 | 48 | Breast | Invasive carcinoma of no special type | 2 | IIB | Malignant | Fmg140095 | +++ | +++ | 0 | +, 10% |
| G6 | 2 | 102 | 55 | Breast | Invasive lobular carcinoma | - | IIIA | Malignant | Fmg100027 | + | - | 3+ | - |
| C10 | 2.75 | 42 | 50 | Breast | Invasive carcinoma of no special type | 2 | IIIB | Malignant | Fmg010912 | +++ | +++ | 0 | +, 5% |
| C12 | 2 | 44 | 52 | Breast | Invasive carcinoma of no special type | 2 | IIB | Malignant | Fmg020652 | +++ | - | 3+ | +, 20% |
| C13 | 1 | 45 | 47 | Breast | Invasive carcinoma of no special type | 2 | IIB | Malignant | Fmg030518 | + | + | 3+ | +, 10% |
| A10 | | 10 | 56 | Lymph node | Metastatic carcinoma from breast | 2 | - | Metastasis | Fmg020026 | + | - | 0 | +, 15% |
| B10 | 2.5 | 26 | 49 | Lymph node | Metastatic carcinoma from breast | 2 | - | Metastasis | Fmg080061 | +++ | - | 0 | +, 65% |
| B8 | 1.5 | 24 | 62 | Lymph node | Metastatic carcinoma from breast | 3 | - | Metastasis | Fmg060484 | + | - | 0 | +, 15% |
| I14 | 1 | 142 | 55 | Breast | Ductal carcinoma in situ | - | 0 | Malignant | Fmg032009 | - | - | 3+ | +, 5% |
| D8 | 0 | 56 | 63 | Breast | Invasive carcinoma of no special type | 1 | IIB | Malignant | Fmg140039 | +++ | - | 2+ | +, 1% |
| E8 | 2 | 72 | 38 | Breast | Invasive carcinoma of no special type | 2 | IIB | Malignant | Fmg010758 | - | - | 3+ | +, 5% |
| G10 | 2 | 106 | 54 | Breast | Invasive lobular carcinoma | - | IIB | Malignant | Fmg100044 | ++ | + | 0 | - |
| H2 | 2 | 114 | 40 | Breast | Invasive lobular carcinoma | - | IIB | Malignant | Fmg060878 | ++ | + | 0 | +, 3% |
| H15 | 3 | 127 | 36 | Breast | Ductal carcinoma in situ with invasion | * | 0 | Malignant | Fmg050788 | +++ | +++ | 0 | +, 5% |
| E10 | 2.5 | 74 | 55 | Breast | Invasive carcinoma of no special type | 2 | IIIB | Malignant | Fmg010908 | ++ | - | 3+ | +, 5% |
| G7 | 0.5 | 103 | 45 | Breast | Invasive lobular carcinoma | - | IIA | Malignant | Fmg120101 | +++ | +++ | 0 | +, 3% |
| A15 | | 15 | 38 | Lymph node | Metastatic carcinoma from breast | 2 | - | Metastasis | Fmg100103 | - | - | 0 | +, 10% |
| B12 | | 28 | 54 | Lymph node | Metastatic carcinoma from breast | 2 | - | Metastasis | Fmg100101 | ++ | - | 2+ | +, 3% |
| H16 | 1 | 128 | 62 | Breast | Ductal carcinoma in situ | - | 0 | Malignant | Fmg130097 | - | - | 3+ | +, 5% |
| D12 | 1.5 | 60 | 53 | Breast | Invasive carcinoma of no special type | 2 | IIA | Malignant | Fmg010551 | - | - | 3+ | +, 20% |
| D10 | 1 | 58 | 32 | Breast | Invasive carcinoma of no special type | 2 | IIA | Malignant | Fmg010731 | +++ | +++ | 0 | +, 5% |
| G2 | 0.5 | 98 | 48 | Breast | Invasive carcinoma of no special type | 1 | IIIA | Malignant | Fmg110129 | - | - | 3+ | - |
| A6 | 2.5 | 6 | 56 | Lymph node | Metastatic carcinoma from breast | 2 | - | Metastasis | Fmg020431 | - | - | 3+ | +, 5% |

| | | | | | | | | | | | | | |
|---|---|---|---|---|---|---|---|---|---|---|---|---|---|
| B5 | 0.5 | 21 | 42 | Lymph node | Metastatic carcinoma from breast | 3 | - | Metastasis | Fmg010821 | ++ | + | 0 | - |
| G9 | 2 | 105 | 69 | Breast | Invasive lobular carcinoma | - | IIA | Malignant | Fmg020930 | + | - | 0 | - |
| G3 | 2.5 | 99 | 42 | Breast | Invasive carcinoma of no special type | 1 | IIB | Malignant | Fmg110138 | +++ | - | 2+ | +, 8% |
| F11 | 2 | 91 | 46 | Breast | Invasive carcinoma of no special type | 3 | IIIA | Malignant | Fmg140115 | - | - | 2+ | +, 3% |
| A14 | 1.5 | 14 | 42 | Lymph node | Metastatic carcinoma from breast (sparse) | - | - | Metastasis | Fmg020539 | - | - | 3+ | +, 5% |
| G15 | 0.25 | 111 | 47 | Breast | Invasive lobular carcinoma | - | IIIB | Malignant | Fmg032391 | +++ | ++ | 0 | +, 8% |
| K1 | 2.5 | 161 | 37 | Breast | Ductal papillomatosis with hyperplasia | - | - | Hyperplasia | Fmg010052 | ++ | + | 0 | +, 3% |
| I16 | 0 | 144 | 51 | Breast | Ductal carcinoma in situ | - | 0 | Malignant | Fmg010539 | +++ | +++ | 0 | - |
| C11 | 2 | 43 | 62 | Breast | Invasive carcinoma of no special type | 2 | IIB | Malignant | Fmg010351 | - | - | 3+ | +, 1% |
| A12 | 0.25 | 12 | 40 | Lymph node | Metastatic carcinoma from breast | 2 | - | Metastasis | Fmg120124 | +++ | ++ | 1+ | +, 10% |
| F14 | 2 | 94 | 50 | Breast | Invasive carcinoma of no special type | 3 | IIIA | Malignant | Fmg110025 | - | - | 2+ | +, 55% |
| C6 | 2 | 38 | 64 | Breast | Invasive carcinoma of no special type (sparse) | 1 | IIA | Malignant | Fmg010825 | +++ | ++ | 0 | +, 3% |
| E16 | 0 | 80 | 45 | Breast | Invasive carcinoma of no special type | 1 | IIA | Malignant | Fmg010760 | +++ | +++ | 0 | +, 30% |
| G13 | 1 | 109 | 38 | Breast | Invasive lobular carcinoma | - | IIA | Malignant | Fmg020138 | +++ | +++ | 0 | +, 10% |
| H13 | 2 | 125 | 44 | Breast | Squamous cell carcinoma | 2 | IB | Malignant | Fmg090046 | - | - | 0 | +, 10% |
| G11 | 0.25 | 107 | 59 | Breast | Invasive lobular carcinoma | - | IIIB | Malignant | Fmg020022 | +++ | +++ | 0 | +, 5% |
| C16 | 0 | 48 | 46 | Breast | Invasive carcinoma of no special type | 2 | IIA | Malignant | Fmg010769 | + | - | 3+ | +, 15% |
| H3 | 3 | 115 | 52 | Breast | Invasive lobular carcinoma | - | IIB | Malignant | Fmg031144 | - | - | 0 | +, 10% |
| J3 | 3 | 147 | 25 | Breast | Fibroadenoma | - | - | Benign | Fmg010046 | +++ | ++ | 0 | - |
| D16 | 2 | 64 | 72 | Breast | Invasive carcinoma of no special type | 2 | IIB | Malignant | Fmg080070 | - | - | 3+ | +, 5% |
| A8 | | 8 | 55 | Lymph node | Metastatic carcinoma from breast | 2 | - | Metastasis | Fmg010941 | - | - | 0 | +, 3% |
| E13 | 0 | 77 | 45 | Breast | Invasive carcinoma of no special type | 2 | IIA | Malignant | Fmg010811 | - | - | 0 | +, 50% |
| C5 | 0.5 | 37 | 30 | Breast | Invasive carcinoma of no special type | 2 | IIIA | Malignant | Fmg090030 | - | - | 3+ | +, 20% |
| F16 | 0.5 | 96 | 45 | Breast | Invasive carcinoma of no special type | 3 | IIB | Malignant | Fmg110098 | +++ | - | 3+ | +, 5% |
| B4 | 0.5 | 20 | 58 | Lymph node | Metastatic carcinoma from breast | 2 | - | Metastasis | Fmg010766 | +++ | +++ | 0 | +, 3% |
| I10 | 0.5 | 138 | 63 | Breast | Ductal carcinoma in situ | - | 0 | Malignant | Fmg110044 | +++ | +++ | 2+ | +, 5% |
| F6 | 2.5 | 86 | 45 | Breast | Invasive carcinoma of no special type | * | IIA | Malignant | Fmg110141 | - | - | 0 | +, 15% |
| H14 | 2.5 | 126 | 47 | Breast | Squamous cell carcinoma | 3 | IA | Malignant | Fmg010930 | - | - | 3+ | +, 5% |
| C9 | 2.5 | 41 | 67 | Breast | Invasive carcinoma of no special type | 1 | IIA | Malignant | Fmg010624 | +++ | - | 0 | +, 5% |
| D11 | 2 | 59 | 52 | Breast | Invasive carcinoma of no special type | 2 | IIIB | Malignant | Fmg020350 | +++ | ++ | 0 | - |
| H10 | 1 | 122 | 45 | Breast | Invasive lobular carcinoma | - | IIB | Malignant | Fmg100047 | +++ | - | 2+ | +, 15% |
| D14 | 2 | 62 | 44 | Breast | Invasive carcinoma of no special type | 2 | IA | Malignant | Fmg010733 | +++ | +++ | 0 | +, 3% |
| F3 | 0.5 | 83 | 71 | Breast | Invasive carcinoma of no special type | 3 | IIIA | Malignant | Fmg020095 | +++ | ++ | 0 | +, 5% |
| F7 | 2 | 87 | 38 | Breast | Invasive carcinoma of no special type | 3 | IIA | Malignant | Fmg010860 | +++ | +++ | 0 | +, 10% |
| J6 | 2 | 150 | 27 | Breast | Adenosis | - | - | Inflammation | Fmg010028 | ++ | ++ | 0 | +, 8% |
| L4 | 1.5 | 180 | 42 | Breast | Cancer adjacent breast tissue | * | - | AT | Fmg020150 | + | + | 0 | - |
| J1 | 1.5 | 145 | 36 | Breast | Fibroadenoma | - | - | Benign | Fmg090075 | + | ++ | 0 | - |
| D9 | 2 | 57 | 45 | Breast | Invasive carcinoma of no special type | 2 | IA | Malignant | Fmg010547 | - | - | 0 | +, 1% |
| E5 | 1 | 69 | 38 | Breast | Invasive carcinoma of no special type | 2 | IIA | Malignant | Fmg010910 | + | ++ | 0 | +, 10% |
| J13 | 2.5 | 157 | 48 | Breast | Mild atypical hyperplasia of duct | - | - | Hyperplasia | Fmg010508 | +++ | ++ | 0 | - |
| K7 | 2.5 | 167 | 29 | Breast | Plasma cell mastitis | - | - | Inflammation | Fmg021965 | * | * | * | * |
| I5 | 1.5 | 133 | 46 | Breast | Ductal carcinoma in situ | - | 0 | Malignant | Fmg031516 | +++ | +++ | 0 | +, 5% |
| E4 | 2 | 68 | 65 | Breast | Invasive carcinoma of no special type | 2 | IIIA | Malignant | Fmg010189 | - | - | 0 | +, 10% |
| B9 | 1.25 | 25 | 39 | Lymph node | Metastatic carcinoma from breast | 2 | - | Metastasis | Fmg080063 | +++ | +++ | 2+ | - |
| D7 | 1.5 | 55 | 44 | Breast | Invasive carcinoma of no special type | 1 | IIA | Malignant | Fmg010859 | +++ | +++ | 0 | +, 10% |
| F9 | 0.25 | 89 | 47 | Breast | Invasive carcinoma of no special type | 3 | IIB | Malignant | Fmg110037 | ++ | +++ | 2+ | +, 50% |

| | | | | | | | | | | | | | |
|---|---|---|---|---|---|---|---|---|---|---|---|---|---|
| K6 | | 166 | 39 | Breast | Chronic mastitis No.135 | * | - | Inflammation | Fmg030989 | ++ | ++ | 0 | - |
| C8 | 1 | 40 | 30 | Breast | Invasive carcinoma of no special type | 1 | IA | Malignant | Fmg020080 | +++ | +++ | 0 | +, 3% |
| E12 | 0.75 | 76 | 43 | Breast | Invasive carcinoma of no special type | 2 | IIA | Malignant | Fmg010835 | +++ | +++ | 0 | +, 5% |
| E14 | 2 | 78 | 45 | Breast | Invasive carcinoma of no special type | 2 | IIIA | Malignant | Fmg130001 | +++ | + | 3+ | +, 5% |
| J10 | 2.5 | 154 | 45 | Breast | Adenosis | - | - | Inflammation | Fmg060022 | ++ | ++ | 0 | - |
| J5 | 1 | 149 | 44 | Breast | Adenosis | - | - | Inflammation | Fmg010032 | ++ | ++ | 0 | - |
| L8 | 2 | 184 | 35 | Breast | Adjacent normal breast tissue | - | - | NAT | Fmg010800 | + | + | 0 | - |
| L10 | 2.5 | 186 | 45 | Breast | Cancer adjacent breast tissue | - | - | AT | Fmg021586 | + | + | 0 | - |
| K15 | 1.75 | 175 | 49 | Breast | Cancer adjacent breast tissue (chronic mastitis) | - | - | AT | Fmg050296 | + | + | 0 | - |
| K11 | 1.5 | 171 | 44 | Breast | Chronic mastitis | - | - | Inflammation | Fmg040540 | + | + | 0 | - |
| J2 | 1 | 146 | 34 | Breast | Fibroadenoma | - | - | Benign | Fmg031932 | +++ | ++ | 0 | - |
| J4 | 2.5 | 148 | 23 | Breast | Fibroadenoma | - | - | Benign | Fmg010048 | +++ | ++ | 0 | - |
| J12 | 3 | 156 | 46 | Breast | Adjacent normal breast tissue (sparse) | - | - | NAT | Fmg060211 | ++ | ++ | 0 | - |
| L7 | 2 | 183 | 46 | Breast | Cancer adjacent breast tissue | - | - | AT | Fmg030942 | + | + | 0 | +, 1% |
| J9 | 3 | 153 | 19 | Breast | Fibroadenoma | - | - | benign | Fmg010034 | ++ | ++ | 0 | +, 1% |
| D13 | 0 | 61 | 42 | Breast | Invasive carcinoma of no special type | 2 | IIA | Malignant | Fmg010917 | +++ | +++ | 0 | +, 3% |
| J15 | 1 | 159 | 38 | Breast | Hyperplasia | - | - | Hyperplasia | Fmg060137 | ++ | ++ | 0 | - |
| J7 | 2.5 | 151 | 23 | Breast | Fibroadenoma | - | - | Benign | Fmg010045 | +++ | ++ | 0 | +, 1% |
| K8 | 2 | 168 | 42 | Breast | Acute mastitis | - | - | Inflammation | Fmg020849 | ++ | - | 0 | - |
| L11 | 2 | 187 | 55 | Breast | Adjacent normal breast tissue | - | - | NAT | Fmg130013 | + | + | 0 | - |
| K14 | 1.75 | 174 | 53 | Breast | Cancer adjacent breast tissue | * | - | AT | Fmg140096 | + | + | 0 | +, 3% |
| L6 | 2.5 | 182 | 57 | Breast | Cancer adjacent breast tissue | * | - | AT | Fmg021442 | + | + | 0 | - |
| K13 | 1.5 | 173 | 29 | Breast | Chronic mastitis of No.66 | - | - | Inflammation | Fmg140093 | + | + | 0 | +, 1% |
| J8 | 3 | 152 | 45 | Breast | Fibroadenoma | - | - | Benign | Fmg060126 | ++ | +++ | 0 | +, 10% |
| J14 | 1 | 158 | 82 | Breast | Hyperplasia | - | - | Hyperplasia | Fmg060093 | + | ++ | 0 | - |
| C14 | 2 | 46 | 43 | Breast | Invasive carcinoma of no special type | * | IA | Malignant | Fmg010839 | +++ | +++ | 0 | +, 3% |
| D15 | 0.25 | 63 | 48 | Breast | Invasive carcinoma of no special type | 2 | IIA | Malignant | Fmg110079 | +++ | - | 2+ | +, 3% |
| J16 | 1.5 | 160 | 40 | Breast | Mild atypical hyperplasia of duct | - | - | Hyperplasia | Fmg060063 | +++ | +++ | 0 | +, 1% |
| K5 | 1.5 | 165 | 39 | Breast | Plasma cell mastitis | - | - | Inflammation | Fmg010013 | ++ | ++ | 0 | - |

**Table S1: Scoring breakdown of emerin staining in tissue microarray samples Individual scores of tissues analyzed in A) figure 9A-B and B) figure 9C-D. N=5 fields of grading per tissue.**
