## Supplemental Figure 1 for "Emerin deficiency drives MCF7 cells to an invasive phenotype"

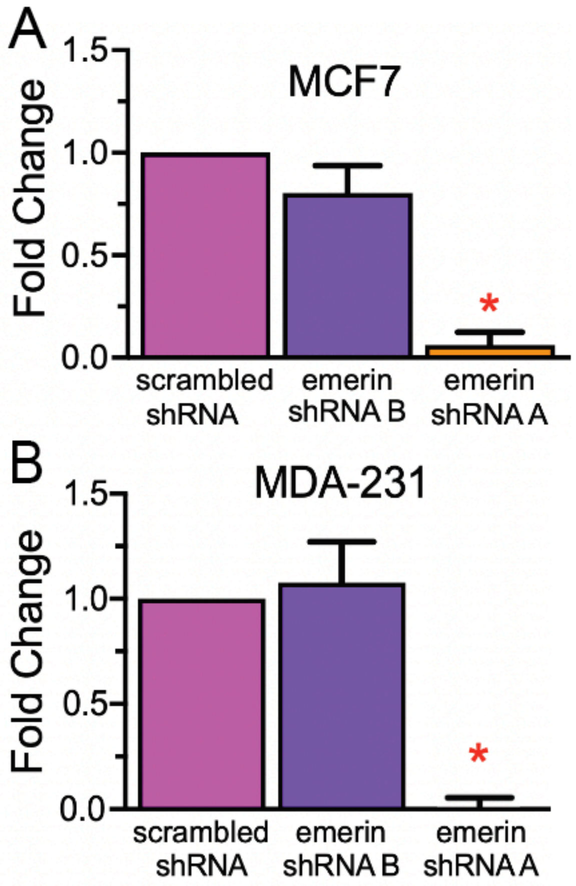


**Figure S1:** Emerin mRNA levels of scrambled shRNA, emerin shRNA B, and emerin shRNA A in A) MCF7 (N=4) and B) MDA-231 (N=3) cell lines, as determined by RT-qPCR. Significance was determined by one-way ANOVA followed by Dunnett's test. *P<0.0001, compared to scrambled shRNA and emerin shRNA B.
