## Supplemental Figure 2 for "Emerin deficiency drives MCF7 cells to an invasive phenotype"

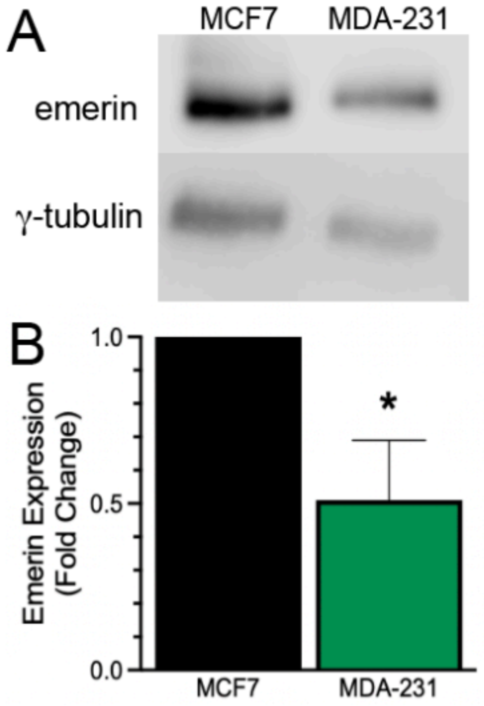


**Figure S2:** Relative emerin levels in poorly invasive MCF7 and highly invasive MDA-231 cells. A) Representative western blot and B) quantification of emerin protein in MCF7 and MDA-231 cell lines normalized to γ-tubulin. MDA-231 cells have 49% less emerin expression than poorly invasive MCF7 cells; N=4. *P=0.0342, Student's t-test.
